# *Paraburkholderia* endosymbionts with streamlined genomes are disadvantaged during interspecific competition

**DOI:** 10.64898/2026.11.24.769019

**Authors:** Calum J. Stephenson, Elle Belperio, David C. Queller, Joan E. Strassmann

## Abstract

Trait loss and genome size reduction are classic evolutionary signatures of obligate bacterial endosymbiosis. However, many bacteria are facultative endosymbionts and so experience a trade-off between adaptation to living within their hosts and apart from them. A central tension in this transition is whether traits required for independent life, including adaptations for interspecific competition, are maintained or lost. We compared three facultatively endosymbiotic *Paraburkholderia* species associated with the social amoeba *Dictyostelium discoideum* to test whether host adaptation is associated with reduced competitive ability. Two species, *P. bonniea* and *P. hayleyella*, display relatively small, streamlined genomes, while the other, *P. agricolaris*, has an unreduced genome compared to free-living congeners. We competed these species and three non-amoeba-associated *Paraburkholderia* against six ecologically relevant competitor bacteria, using fluorescent markers to track strain-specific fitness in co-culture. The reduced-genome species were never more competitive than *P. agricolaris* and, unlike *P. agricolaris,* never outcompeted their competitors. This deficit was reflected in a reduced ability to grow quickly to high levels in co-culture. Follow-up assays revealed that cell-free supernatant from the co-culture of *P. agricolaris* and a *Pseudomonas* isolate inhibited subsequent growth of the *Pseudomonas*, whereas supernatant from the co-culture of *P. hayleyella* and *Pseudomonas* did not. Reduced genome size in these facultative endosymbionts is associated with weaker competitive performance without eliminating independent proliferation under laboratory conditions. This competitive erosion may lead to ecological host dependence, where symbionts remain physiologically capable of living independently but become increasingly unable to persist outside the host in competitive microbial communities.

## Introduction

Competition over limited resources such as nutrients and space is a fundamental force on the evolution of bacteria in nature [1–3]. This pressure has driven the evolution of many traits for both offence and defence, from metabolic flexibility to toxin-antitoxin systems, siderophore production, biofilm formation, secretion systems, and motility [4–6]. Some bacteria evade competitive interactions by escaping inside larger, typically eukaryotic cells. If these bacteria can avoid digestion and the host immune system, they can enjoy freedom from many of their natural abiotic and biotic stressors, and often have access to nutrients and biosynthetic intermediates within their hosts. When bacteria routinely spend periods of their life cycles inside host cells, they are called endosymbionts.

The evolution of endosymbiosis has attracted significant research attention, partly due to its relevance to the understanding of mutualism and pathogenesis [7–10]. Both empirical and theoretical research suggest a common mechanism where the simple intracellular environment relaxes selection for bacterial traits key for extracellular survival, leading to gene loss through genome streamlining [11,12]. In obligately host-dependent symbionts, this can be compounded by a smaller effective population size in the intracellular environment. This increases the relative strength of genetic drift and causes the loss of more genetic material through mutational degradation and pseudogenisation [13–15]. These processes often result in the so-called “endosymbiotic syndrome”: small, AT-rich genomes [16], reduced metabolic capacity [13,17,18], decreased tolerance of environmental stress [19,20], and reduced defences against phage predation and interbacterial competition [21].

However, much of the experimental work underlying these observations has been focused on highly derived, obligate endosymbionts that are unable to live without their hosts. Many bacterial endosymbionts are facultatively host–associated and retain the ability to live outside host cells [22,23]. These bacteria may therefore face a selective trade-off between the benefits of streamlining in host-associated environments and the costs of losing traits required for extracellular survival. For endosymbionts, one of the biggest differences between intracellular and extracellular environments is the diversity of the surrounding bacterial community and the intensity of interspecific competition.

In this study, we explore how interspecific competitive ability is shaped by a history of genome streamlining during the transition to a facultative endosymbiotic lifestyle within the bacterial genus *Paraburkholderia*. *Paraburkholderia* are soil-dwelling betaproteobacteria, often found in association with plants [24]. However, three species of *Paraburkholderia* are known to readily form endosymbiotic associations with the social amoeba *Dictyostelium discoideum* [25], and can also do so in the lab with a number of other Dictyostelidae species [26]. Named *Paraburkholderia agricolaris*, *P. bonniea*, and *P. hayleyella* [27], they represent two phylogenetically independent transitions towards a *D. discoideum*-associated existence [25,28]. *P. hayleyella* and *P. bonniea* are each other’s closest relatives and appear to have an extensive history as *D. discoideum* endosymbionts, with their 4.1 Mbp genomes being the smallest within *Paraburkholderia* [29] and around 50% smaller than the genus median. This has led to a limited capacity for metabolising different carbon sources and narrower abiotic tolerances for these two species [27]. This reduction appears to be mostly a product of genome streamlining rather than degradation, based on pseudogene content and gene density [30]. By contrast, *P. agricolaris* emerged within a different clade and appears to have transitioned towards facultative endosymbiosis more recently, with a genome typical of free–living *Paraburkholderia* in terms of both size and content [27,30]. Furthermore, *P. hayleyella* achieves greater fitness as measured by population sizes in the presence of *D. discoideum* [31,32]; *P. agricolaris* saw no fitness change with or without *D. discoideum,* while *P. bonniea* was not examined for this trait.

Despite these differences, there are many traits shared among these species. All three are ostensibly facultative symbionts, as they can be readily cultivated without their hosts under laboratory conditions. All three reduce host spore production [33,34] but provide context-dependent benefits upon dispersal to food-poor environments through the induction of secondary bacterial carriage [35,36]. And all three also possess a homologous genomic island encoding both a Type III and Type VI secretion system, absent from other *Paraburkholderia*, with an apparent role in host infection [30,34,37].

To assess whether the transition towards facultative endosymbiosis has shaped competitive fitness in this group, we competed four strains from each of the three symbiotic *Paraburkholderia* species against six species of free-living soil bacteria that are edible to *D. discoideum*. Additionally, we competed the same set of soil bacteria against three species of *Paraburkholderia* without a known history of association with *D. discoideum*. To independently track the growth of each species in co-culture, we differentially tagged each species with fluorescent proteins [38]. This allowed us to quantify the relative competitive performance of both bacterial partners and investigate whether competitive outcomes were associated with changes in fluorescence yield and maximum growth rate. If increased association with *D. discoideum* has relaxed selection for interbacterial competitive fitness, we hypothesise that *P. bonniea* and *P. hayleyella* will exhibit reduced competitive fitness relative to the less symbiotically derived *P. agricolaris*. If amoeba association itself relaxes selection on competitiveness, we also expect that *P. agricolaris* will exhibit reduced competitiveness relative to species not associated with amoebas. However, if interspecies competition remains a strong selective pressure on these species due to their facultative life history, we predict no significant deficit in competitive ability between our species despite the reduced genomes of *P. bonniea* and *P. hayleyella*.

## Materials and Methods

### Isolation of bacterial strains

Earlier we isolated four strains of both *P. agricolaris* (Ba31, Ba80, Ba159, and Ba161) and *P. hayleyella* (Bh11, Bh21, Bh22, and Bh171) as well as two strains of *P. bonniea* (Bb433 and Bb859) from wild strains of *Dictyostelium discoideum* according to previously published protocols [25,28,39]. For this project, we isolated two additional strains of *P. bonniea* derived from *D. discoideum* strains QS822 and QS862, that we name Bb822 and Bb862. We confirmed *P. bonniea* identity using newly designed primers (forward primer 5’-TGATCCCGTGCCCGAATTCACGC-3’ and reverse 5’-ACTCTCCTGGTGGGCAGGCATC-3’) that selectively amplify approximately 500 bp of the *prmA* gene. We also included three strains of Paraburkholderia with no known amoeba association to form our non-amoeba-associated group: *P. silvatlantica* PVA5, *P. unamae* MT1-641 (both received from Ann Hirsch) and *P. fungorum* BAA-463 (Table 1).

**Table 1:**
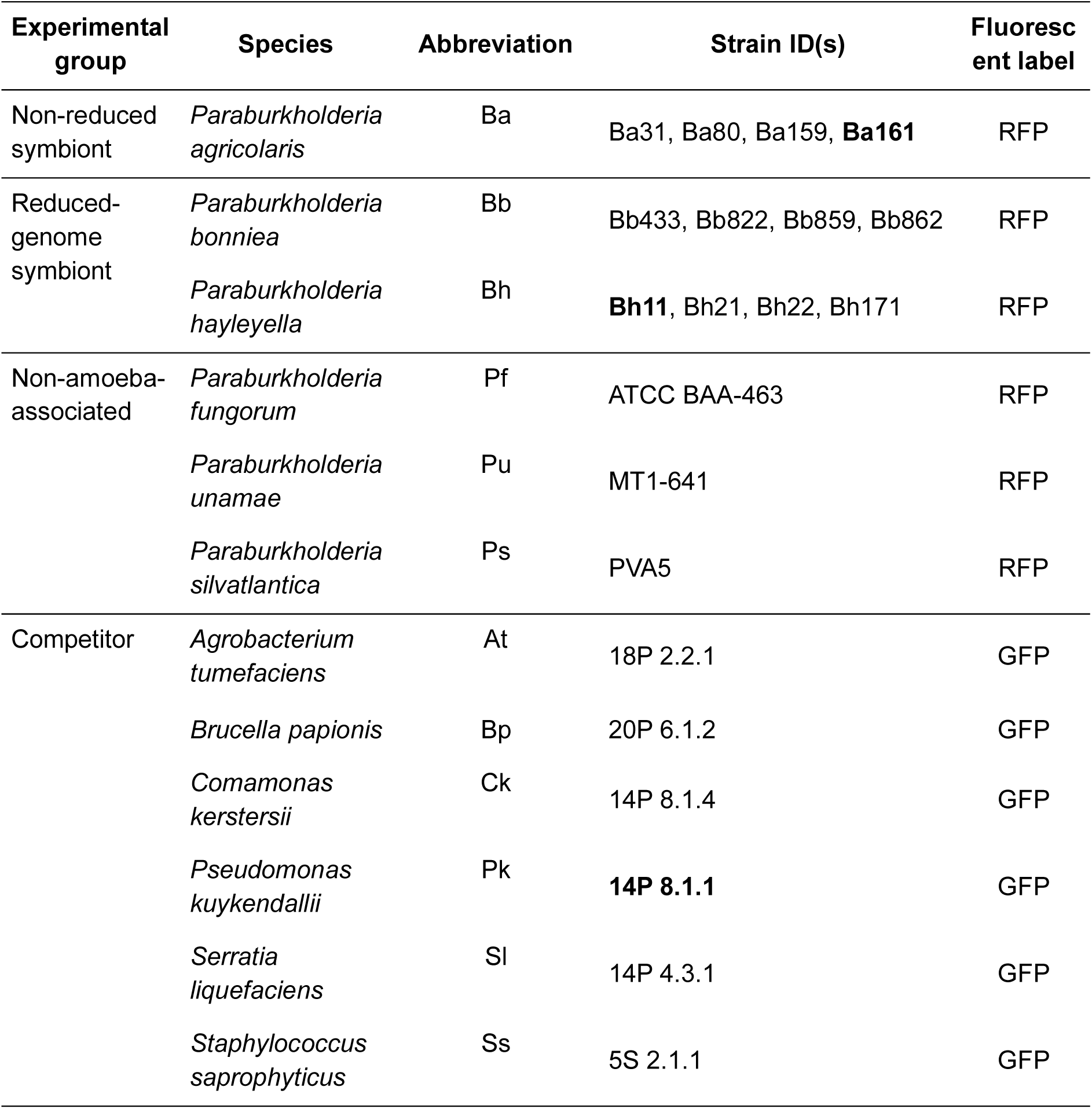
Bacterial strains and fluorescent transconjugants used in competition assays. Species identifications for competitor isolates follow Brock et al. (2018). Strains used in the conditioned-supernatant assays are highlighted in bold. The non-amoeba-associated group comprises three *Paraburkholderia* species with no known association with *Dictyostelium discoideum*.

We used six strains of non-*Paraburkholderia* ‘competitor’ bacteria in this study; *Pseudomonas kuykendallii* 14P 8.1.1 (abbreviated for statistical reporting as Pk), *Agrobacterium tumefaciens* 18P 2.2.1 (At), *Serratia liquefaciens* 14P 4.3.1 (Sl), *Comamonas kerstersii* 14P 8.1.4 (Ck), *Brucella papionis* 20P 6.1.2 (Bp), and *Staphylococcus saprophyticus* 5S 2.1.1 (Ss). We refer to strains using abbreviations for all statistical reporting. All six were isolated from *D. discoideum* fruiting bodies collected from Mountain Lake Biological Station, VA in 2014 [40].

### Fluorescent labelling of bacterial strains

We differentiated *Paraburkholderia* and competitor strains in co-culture using red (RFP) and green (GFP) fluorescent protein labels, respectively. Most of the *Paraburkholderia* strains had been previously tagged by DiSalvo et al. [25], and we repeated the protocol with the plasmids described there to transform the two novel *P. bonniea* isolates in this study. For the competitors, the protocol for GFP introduction was identical except we used the *E. coli* helper MDFpir (pTNS2), donor MDFpir (pUC18R6KT-miniTn7T-Gm-PA1-GFP), and selected for transformants using the antibiotic gentamicin. We confirmed species identity of fluorescent transformants through 16S rRNA sequencing.

### Co-culture experiments

We streaked clonal glycerol stocks onto yeast glucose (YG) agar plates (5 g glucose, 4 g Oxoid yeast extract, 1 g NaCl, 15 g agar per litre ddH_2_0) and incubated them at 30 °C for 24 hours. We then inoculated single colonies into 5 mL of liquid YG and grew cultures overnight at 30 °C while shaking. We sub-cultured the overnight samples 1:10 into fresh YG medium and continued incubation until each strain reached the logarithmic phase of growth. Immediately prior to the experiment, we measured each culture’s optical density at 600 nm (OD_600_) using a BioPhotometer (Eppendorf) and subsequently diluted the culture using fresh media to 0.1 OD_600_.

We established monocultures, fluorescently labelled–unlabelled wild type (WT) controls, and interspecific co-cultures in 96-well plates (Fig. 1A). Each well contained 100 µL of fluorescent bacterial culture (OD_600_ = 0.1) mixed with an equal volume of fresh YG, the corresponding WT strain, or a differently labelled competitor, respectively. Both the unlabelled WT strains and labelled competitor strains were added at an OD_600_ of 0.1. Each treatment was replicated in three different wells on each day. We repeated each set of competition mixtures and labelled-unlabelled WT controls on at least three separate days per strain. All competition and control conditions were paired with a daily monoculture of the fluorescent strains involved. Finally, three wells per day were filled with 200 μL of fresh YG media to act both as a marker for media contamination and as a control for background fluorescence.

**Figure 1:**
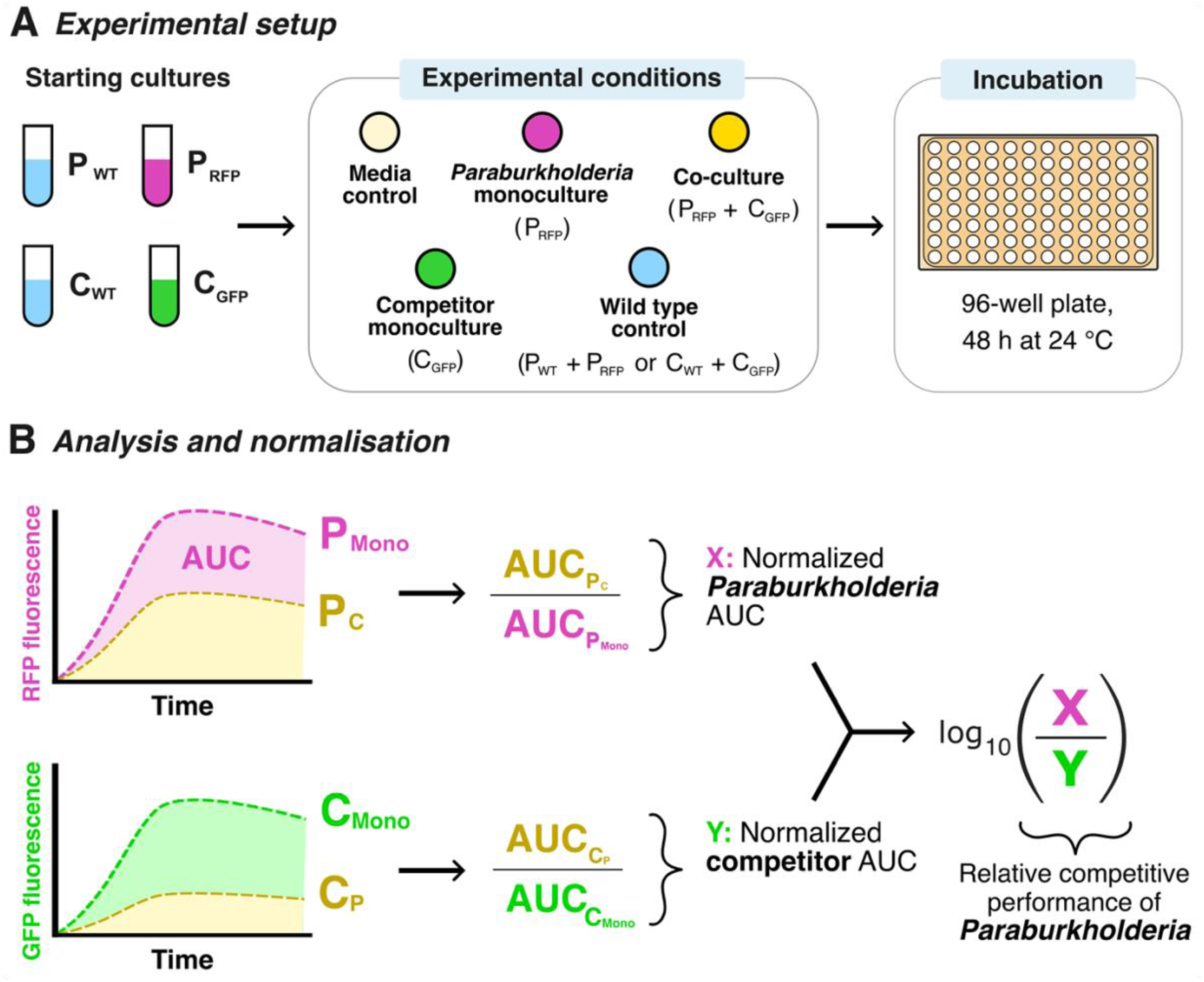
Fluorescence-based assay of bacterial competition. **(A)** Cultures of RFP-expressing *Paraburkholderia* and six different GFP-expressing soil competitors are diluted to OD_600_ = 0.1 in nutrient medium. For each treatment, 100 µL of fluorescently labelled culture was combined in a 96-well plate with either 100 µL of fresh YG (monoculture), a differentially labelled competitor (co-culture), or the corresponding unlabelled wild-type strain (WT control). **(B)** After correcting for media autofluorescence and fluorescence bleed-through between channels, RFP and GFP fluorescence can be used as a proxy for *Paraburkholderia* and competitor growth, respectively. We calculated the area under each fluorescent curve (AUC) in competition and normalised the AUC of each strain against their respective same-day monoculture AUC (X and Y). By taking the log_10_-transformed ratio of X to Y, we calculated the relative competitive performance of *Paraburkholderia* with positive values indicating an advantage to *Paraburkholderia* and negative values indicating an advantage to the competitor.

For all conditions, we grew cultures in a Tecan Spark® microplate reader (Tecan Group Ltd., Switzerland) for 48 hours at 24 °C, using a 96-well plate sealed with a lid. We filled perimeter wells with 200 µL of YG and excluded them from analysis to prevent edge effects. Every hour, cultures were shaken for 5 s before measuring OD_600_, RFP fluorescence (excitation: 555 ± 10 nm; emission: 590 ± 10 nm), and GFP fluorescence (excitation: 485 ± 10 nm; emission: 535 ± 10 nm). Fluorescence measurements were performed using manual gain settings of 70 for RFP and 60 for GFP.

### Starting-ratio and supernatant toxicity assays

Observation of growth dynamics suggested that *P. agricolaris* caused *P. kuykendallii* fluorescence to sharply decline during co-cultures, an effect that was especially prominent when *P. kuykendallii* was grown with *P. agricolaris* strain Ba161. To examine the effects of starting population ratios on these outcomes, we prepared overnight cultures of Ba161 and *P*. *kuykendallii*. We adjusted both cultures to OD_600_ = 0.1 and mixed them at 15 initial Ba161 proportions (0, 0.01, 0.1, 1, 10, 25, 40, 50, 60, 75, 90, 99, 99.9, 99.99, and 100%), with the remaining proportion comprising *P*. *kuykendallii*. We monitored fluorescence over 24 h using the plate-reader procedures described above.

To investigate whether *P. agricolaris* could produce inhibitory supernatants during competition, we prepared monocultures of Ba161 and *P*. *kuykendallii*, as well as *P. hayleyella* strain Bh11 as a control. Following overnight growth, we sub-cultured strains at initial OD_600_ values of 0.1, 0.01, or 0.001 and incubated them for 24 h. We also established co-cultures by combining equal volumes (2.5 mL) of *P*. *kuykendallii* and either *Paraburkholderia* strain, each adjusted to OD_600_ = 0.1, and incubated them for 24 h. We selected cultures with OD_600_ values closest to 1 and collected cell-free supernatants by centrifugation (10,000 × *g*, 5 min) and filtration (0.2 µm).

We tested supernatants from the three monocultures and two co-cultures at a final concentration of 90%, combining 180 µL of supernatant with 20 µL *P*. *kuykendallii* culture to achieve an initial *P*. *kuykendallii* OD_600_ of 0.1. Each treatment was replicated in three wells across six experimental days, alongside bacteria-free supernatant and medium controls. We monitored OD_600_ and fluorescence hourly over 48 h at 24 °C using the Tecan Spark microplate reader, using identical settings to the co-culture competition experiment.

### Data Preparation

We generated separate RFP and GFP growth curves for each well over 48 h, except for mixtures containing *S. saprophyticus*, which were truncated at 36 h due to unusual late-stage biphasic fluorescence patterns. To correct for background fluorescence and fluorescence detected in the unintended channel, we generated strain- and date-specific standard curves using monoculture measurements. Because each monoculture contained only one labelled strain, fluorescence detected in the opposite channel could be used to estimate the unwanted fluorescence contributed by that strain in co-culture. We fitted linear regressions predicting fluorescence in the unintended channel from fluorescence in the primary channel (e.g., RFP fluorescence from GFP fluorescence in GFP-labelled monocultures). We then applied these regressions to the corresponding same-day co-cultures, subtracting the predicted unwanted fluorescence from each channel at every timepoint. This produced corrected fluorescence curves more closely representing the growth of each labelled strain.

We used fluorescence-derived growth curves as a proxy for bacterial performance during interspecific competition [38,41,42]. We calculated the area under the curve (AUC) for each corrected fluorescence growth curve using the R package *MESS*. To normalise competitive performance across strains and experimental dates, we divided each co-culture AUC by the mean AUC of the corresponding fluorescent bacterial strain grown in monoculture on the same day (Fig. 1B). We calculated relative *Paraburkholderia* competitive performance as the ratio of normalised RFP AUC (*Paraburkholderia*) to normalised GFP AUC (competitor). This ratio was log₁₀-transformed for statistical analysis.

We estimated fluorescence yield (*K*) over 0–36 h as the 98th percentile of LOESS-smoothed fluorescence curves meeting predefined logistic-fit quality criteria. Otherwise, we used the maximum fluorescence before a sustained decline, or the mean of the final 10% of readings when no decline occurred. We calculated μ_max_ over 0–24 h as the steepest five-point rolling slope of log-transformed fluorescence, retaining windows only when at least three readings exceeded the baseline noise threshold (median 0–2 h fluorescence plus two median absolute deviations). Both traits were normalised to same-day monocultures and expressed as log₂ fold changes relative to the median labelled–unlabelled WT control.

### Statistical Analyses

All statistical analyses were performed in R (version 4.3.2) [43]. We averaged technical replicates within each strain–competitor combination and experimental date. We analysed the log₁₀-transformed ratio of monoculture-normalised RFP to GFP AUC using Gaussian mixed-effects models in *glmmTMB*. *Paraburkholderia* species, competitor identity, and their interaction were included as fixed effects, and experimental date and *Paraburkholderia* strain as random intercepts. We compared candidate residual-variance structures using maximum likelihood (ML), with AIC and BIC both favouring the same model. We refitted the selected model using restricted maximum likelihood (REML) for inference. The final model allowed residual variance to vary additively with *Paraburkholderia* species and competitor identity (Table S3).

We calculated estimated marginal means using *emmeans*. Planned comparisons included six pairwise contrasts among the four *Paraburkholderia* groups within each competitor (Holm correction within competitor) and 24 tests against competitive parity (log₁₀ ratio = 0; Holm correction across all tests). We also compared the four non-reduced species with the two reduced species, and the three non-amoeba-associated species with the two reduced species. Each species contributed equally to its respective group mean, with contrasts averaged equally across competitors and Holm correction across the two pooled comparisons. A robust mixed-model sensitivity analysis of *P. agricolaris* Ba161 did not alter model predictions.

To examine absolute deviations from monoculture, we separately modelled log₁₀-normalised RFP and GFP AUC using the same model structure. Overall estimated marginal means were averaged equally across all six *Paraburkholderia* species and six competitors, back-transformed to percentage changes, and tested against zero using asymptotic Wald tests with Holm correction across the two responses (Fig. S2B).

We analysed fluorescence-derived *K* and μ_max_ separately for RFP and GFP. Candidate Gaussian mixed-effects models included *Paraburkholderia* species, competitor identity, and their interaction as fixed effects, with alternative random-effect and residual-variance structures. We selected models based on AIC under ML, with BIC also reported for comparison, and refitted the selected models using REML (Table S6). We used *emmeans* to test deviations from WT-control baselines and compare the four *Paraburkholderia* groups, both averaged across competitors and within individual competitors, applying Holm correction within each test family. The selected models differed in whether they included experimental date alone or date plus *Paraburkholderia* strain as random effects, while all four allowed residual variance to vary additively with *Paraburkholderia* species and competitor identity (Table S6).

For conditioned-supernatant assays, we analysed log₁₀-transformed GFP AUC normalised to same-day *Pseudomonas* monoculture controls grown in fresh YG using a linear model, with conditioned-supernatant treatment and experimental date as fixed effects. We used *emmeans* for planned comparisons between treatments and tests of treatment means against zero, applying Holm correction within each test family (Table S8).

## Results

### Competition suppressed bacterial growth relative to growth alone

We competed each strain of *Paraburkholderia* against six species of free-living soil bacteria at equal 1:1 starting ratios in nutrient-rich YG medium and measured the relative fitness of both species during co-culture (Fig. 1). As expected, co-culture generally reduced performance relative to monoculture, with model-estimated reductions in AUC of 57.0% for *Paraburkholderia* (95% CI: 54.2–59.6%) and 30.8% for competitors (95% CI: 27.9–33.7%; both Holm-adjusted p < 0.001; Fig. S2B). However, the magnitude of these reductions varied among bacterial combinations. Only *Staphylococcus* grew as well in competition as it did alone when pitted against all *Paraburkholderia* except *P. agricolaris* (Fig. S2A).

### Competitive performance varies markedly among *Paraburkholderia* groups

*P. agricolaris* (Ba) exhibited a significantly higher competitive fitness relative to *P. bonniea* (Bb) against five of the six competitor species (Ba − Bb significant for At, Bp, Ck, Pk, and Sl; estimates = 0.25 to 0.949, Holm p ≤ 0.046) and was higher than *P. hayleyella* (Bh) for four of the six species (Ba − Bh significant for At, Bp, Pk, and Sl; estimates = 0.175 to 0.849, Holm p ≤ 0.008; Fig. 2; Table S1). Neither of the reduced species was ever more competitive than *P. agricolaris*. Notably, a difference in competitive outcome between the two reduced species was only observed when competing with *Agrobacterium*, against which *P. hayleyella* was more competitively fit than *P. bonniea* (Bb − Bh = −0.364, Holm p ≤ 0.001).

**Figure 2:**
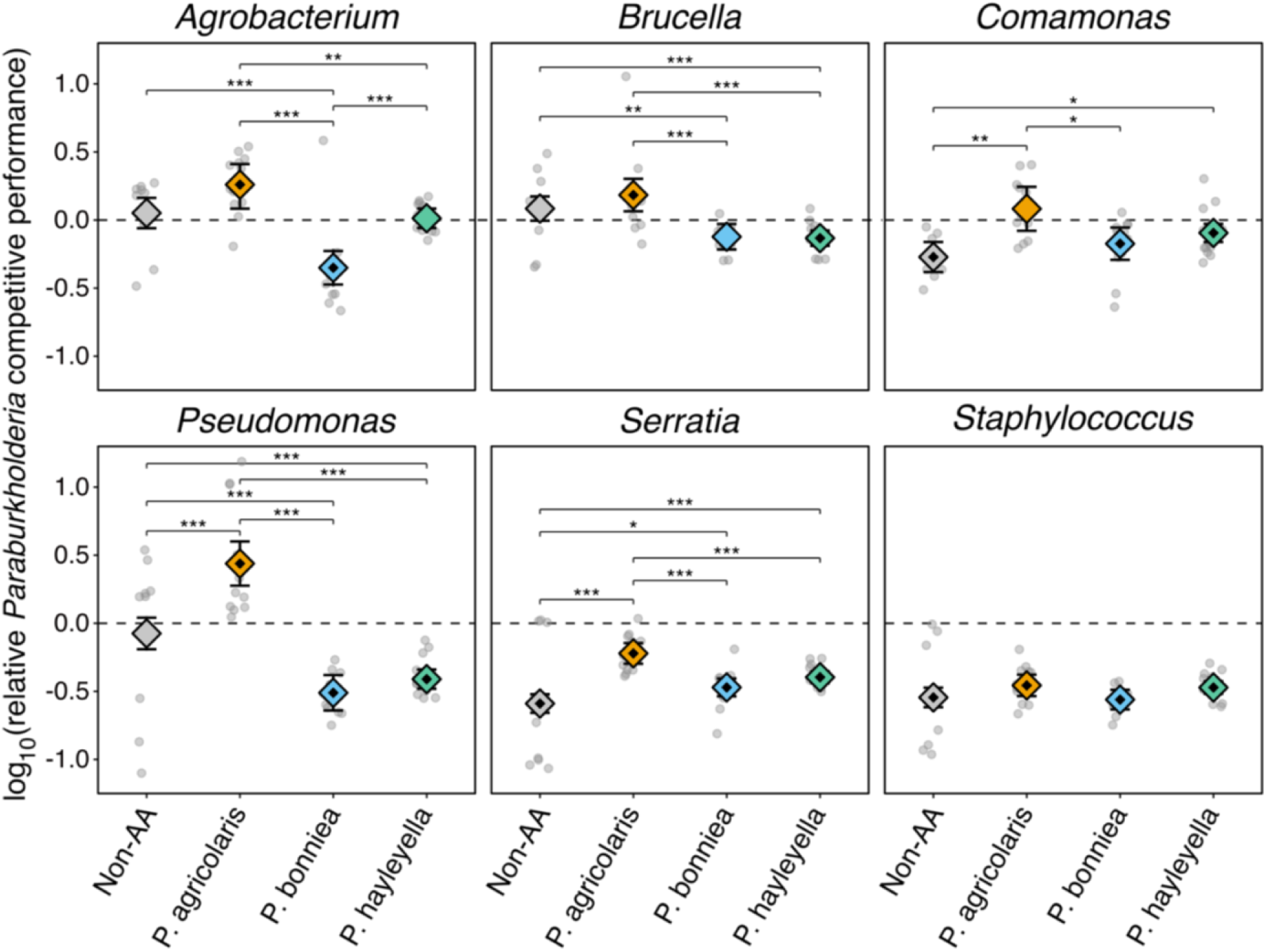
Competition outcomes across competitor strains for each *Paraburkholderia* group. Relative *Paraburkholderia* competitive performance was calculated as the log₁₀-transformed ratio of same-day monoculture-normalised RFP area under the curve (AUC) to the corresponding monoculture-normalised GFP AUC. Values above zero indicate a relative *Paraburkholderia* advantage, whereas values below zero indicate a relative competitor advantage. Grey points represent experimental-date observations, while coloured diamonds show model-estimated marginal means with whiskers indicating 95% confidence intervals. The non-amoeba-associated (Non-AA) group represents the equally weighted mean of three *Paraburkholderia* species. Significance brackets indicate planned pairwise contrasts among *Paraburkholderia* groups, with Holm adjustment across six comparisons within each competitor species (* p < 0.05; ** p < 0.01; *** p < 0.001).

To contextualise these differences, we compared our symbionts with three non-amoeba-associated *Paraburkholderia* species. Of the 12 comparisons, the non-amoeba-associated group outperformed both reduced-genome species in five instances and underperformed them in three. It underperformed relative to the non-reduced symbiont *P. agricolaris* against three of six competitors (Table S1).

### Competitor identity drives co-culture outcomes

Competitor identity (χ² = 760.05, df = 5, p < 0.001) and its interaction with *Paraburkholderia* species (χ² = 500.83, df = 25, p < 0.001) significantly affected competitive outcomes (Fig. 2). Differences among *Paraburkholderia* species were particularly pronounced against *Pseudomonas*, which was outcompeted by *P. agricolaris* but outcompeted both reduced-genome species*. P. agricolaris* was the only *Paraburkholderia* grouping to outperform any competitor, doing so against three of the six strains. In contrast, *P. bonniea* and *P. hayleyella* lost to their competitors in 10 of 12 pairings. The non-amoeba-associated group did not differ significantly from competitive parity against three competitors and lost against *Comamonas, Serratia* and *Staphylococcus*, with the latter two outcompeting all four *Paraburkholderia* groups (Fig. 2; Table S2).

### Genome reduction is associated with lower average competitive performance

Because *P. agricolaris* also exhibited greater competitive performance than the non-amoeba-associated species against several competitors (Fig. 2), we examined whether the apparent competitive disadvantage of the reduced-genome symbionts primarily reflected the unusually strong performance of *P. agricolaris*. We therefore compared the two reduced-genome species with all four non-reduced-genome *Paraburkholderia* species, using model-based contrasts averaged across the six tested competitors. The non-reduced-genome species exhibited significantly greater relative competitive performance (Δlog₁₀ = 0.151, 95% CI: 0.091–0.210, Holm-adjusted p < 0.001; Fig. 3; Table S9), corresponding to a 1.41-fold difference. When the comparison specifically excluded *P. agricolaris*, the difference remained positive and significant, although smaller (Δlog₁₀ = 0.083, 95% CI: 0.010–0.155, Holm-adjusted p = 0.026), corresponding to a 1.21-fold difference.

**Figure 3:**
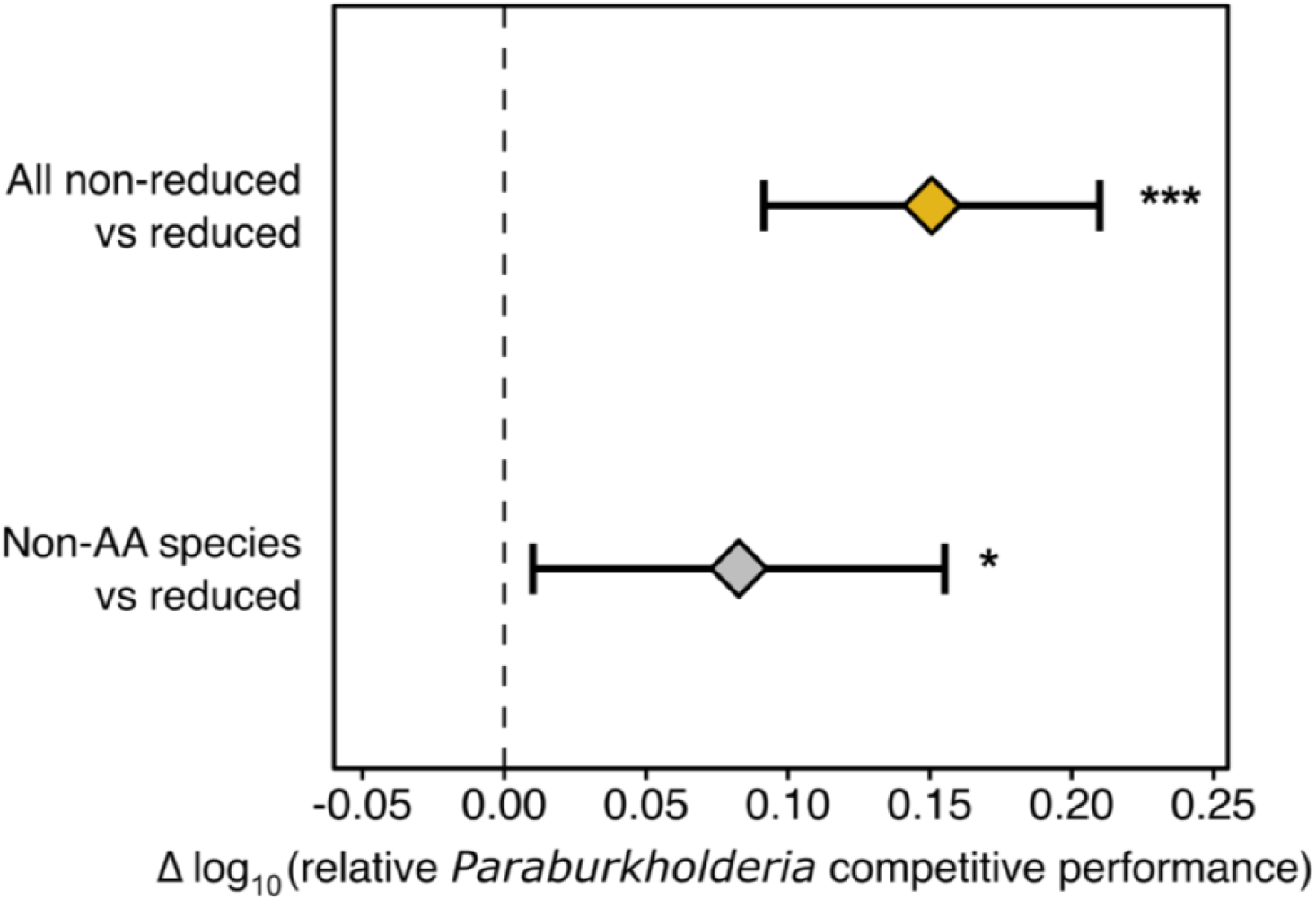
Non-reduced *Paraburkholderia* exhibit greater relative competitive performance than the streamlined lineage, averaged across competitors. Model-based contrasts comparing all four non-reduced-genome species (*P. agricolaris*, *P. fungorum*, *P. unamae* and *P. silvatlantica*; gold diamond) and the three non-amoeba-associated (Non-AA) species (*P. fungorum*, *P. unamae* and *P. silvatlantica*; grey diamond) with the two reduced-genome symbionts (*P. bonniea* and *P. hayleyella*). Contrasts were estimated from the species-by-competitor mixed-effects model, with equal weighting of species within groups and averaging across the six tested competitors. Diamonds represent estimated differences in log₁₀ relative competitive performance between groups, and whiskers indicate 95% confidence intervals. Positive values indicate greater relative competitive performance of the non-reduced-genome or non-amoeba-associated species, respectively, whereas negative values favour the reduced-genome symbionts. Asterisks indicate significance based on Holm-adjusted p values across the two contrasts (* p < 0.05; *** p < 0.001).

### The competitive superiority of *P. agricolaris* is reflected by a robust yield and early growth rate in competition

Following theoretical distinctions between competition acting on carrying capacity and per-capita growth rates [44–46], we examined maximum fluorescence yield (*K*) and maximum growth rate (μ_max_) for both bacterial partners in co-culture, respectively. To capture the specific effect of the competitors on these measures as opposed to the effects of experiencing a higher bacterial concentration, we established a baseline by examining the growth of labelled strains in co-culture with their corresponding unlabelled wild-type (WT) strains (Fig. S4). Reductions in *K* varied around 50% of monoculture, while μ_max_ generally decreased from the 0% expectation (Fig. S4). Growth with WT was therefore used to normalise the growth in competition. Thus, zero indicates no deviation from the empirical WT co-culture baseline.

Neither *P. agricolaris* nor the non-amoeba-associated group differed significantly from their WT-control baseline for *K* and μ_max_ (Figs. 4A, C; Table S5). In contrast, both reduced-genome species showed significant declines in both traits (Bb: *K* = −1.019, μ_max_ = −0.875; Bh: *K* = −0.695, μ_max_ = −0.381; all Holm-adjusted p < 0.001). Comparing across *Paraburkholderia* groups, both reduced-genome species exhibited significantly greater declines relative to *P. agricolaris* in both *K* and μ_max_ (all Holm-adjusted p < 0.001 except Ba–Bh μ_max_, p = 0.021) and *P. bonniea* was additionally weaker than the non-amoeba-associated for both traits (non-AA − Bb: *K* = 0.664, Holm p = 0.008; μ_max_ = 0.814, Holm p < 0.001). Notably, *P. hayleyella* maintained its μ_max_ better than *P. bonniea*, despite both declining from the baseline (Bb − Bh = −0.494, Holm p = 0.008; Fig. 4C, Table S4).

**Figure 4:**
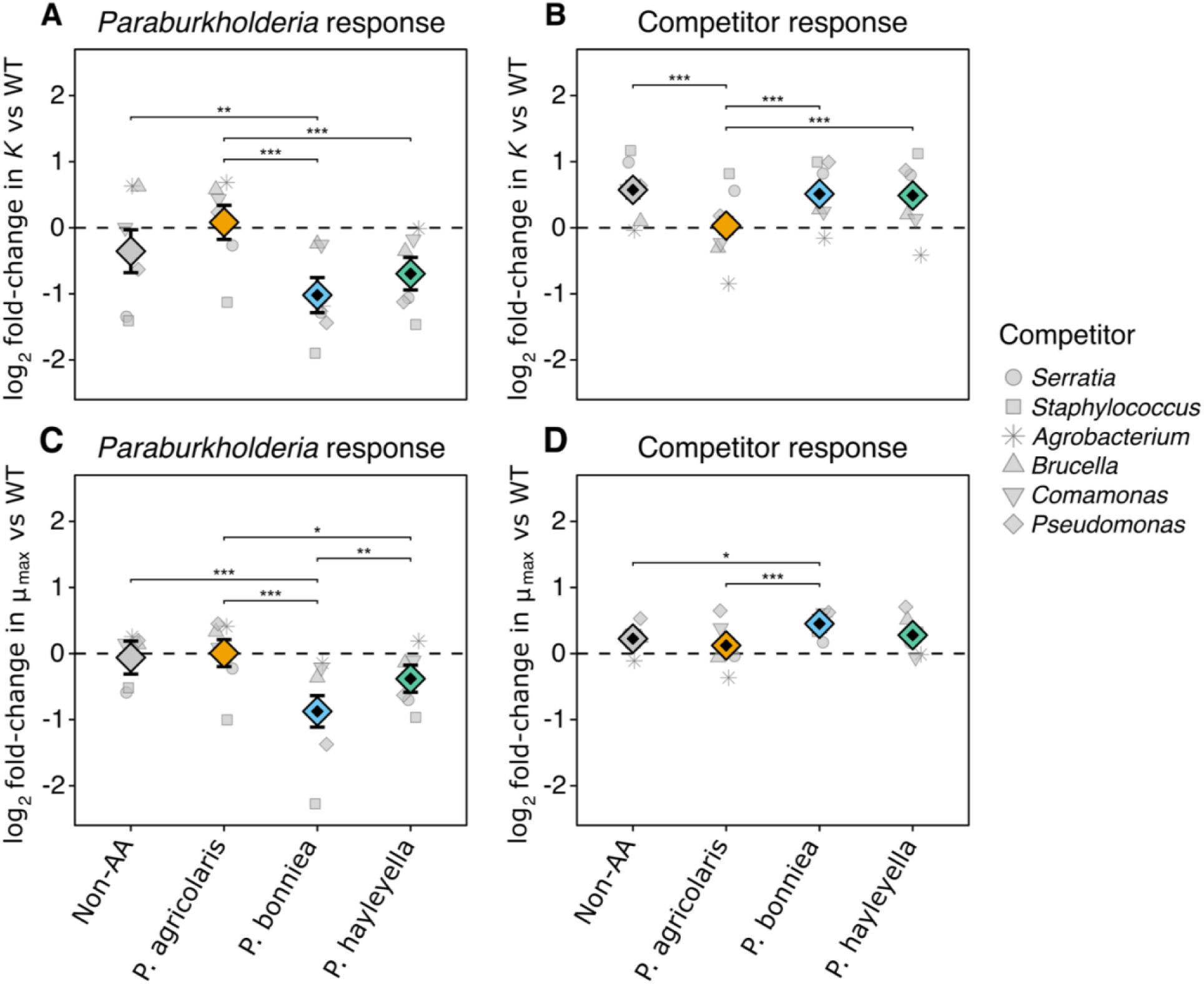
Fluorescence-derived yield (*K*) and maximum growth rate (μ_max_) vary by *Paraburkholderia* identity across both partners in competition. Values are expressed as log₂ fold-changes relative to matched labelled–unlabelled wild-type (WT) co-culture controls, with the dashed line indicating no deviation from this baseline. The left column (A, C) shows the *Paraburkholderia* response, and the right column (B, D) shows the competitor response. The upper row (A, B) shows K, and the lower row (C, D) shows μ_max_. Grey symbols represent competitor-specific mean responses for each *Paraburkholderia* group, with shapes identifying competitor species. Coloured diamonds show model-estimated marginal means, with whiskers indicating 95% confidence intervals. Black centres within diamonds indicate significant departures from the WT baseline, with Holm adjustment across four groups within each response × trait. The non-amoeba-associated (Non-AA) group represents the equally weighted mean of three *Paraburkholderia* species. Significance brackets indicate planned pairwise contrasts among *Paraburkholderia* groups, with Holm adjustment across six comparisons within each response × trait (* p < 0.05; ** p < 0.01; *** p < 0.001).

Competitive ability depends not only on a bacterium’s capacity to maintain its own growth, but also on its effects on the growth of its competitors. Therefore, we also examined how *Paraburkholderia* identity affected competitor traits (Figs. 4B, D; Table S5). On average, competitors exhibited significantly increased μ_max_ when paired with all four *Paraburkholderia* groups, while *K* increased when paired with either reduced-genome species or the non-amoeba-associated group but not with *P. agricolaris* (Figs. 4B, D). Direct comparisons among *Paraburkholderia* groups revealed that competitors paired with *P. agricolaris* exhibited significantly lower *K* than those paired with any other group (Fig. 4B; Table S4). *P. agricolaris* and the non-amoeba-associated also produced a lower competitor μ_max_ than those paired with *P. bonniea* (Fig. 4D). Within-competitor planned contrasts for both partners and both growth traits are presented in the supplementary information (Figs. S5 and S6).

### *P. agricolaris*–*Pseudomonas* co-culture generates a *Pseudomonas*-suppressive supernatant

Competitive outcomes with *Pseudomonas* depended strongly on the *Paraburkholderia* partner (Fig. 2), as it lost to *P. agricolaris* but strongly suppressed both reduced species. Examination of the underlying growth curves revealed strongly non-sigmoidal *Pseudomonas* growth dynamics during *P. agricolaris* co-culture, characterised by rapid early growth followed by a sharp decline in fluorescence after approximately 10 h (Fig. S1). When the starting ratio of the two strains was varied, Ba161 caused sharp declines in *Pseudomonas* growth even when it comprised only 0.1% of the initial population (Fig. 5A; Fig. S7).

**Figure 5:**
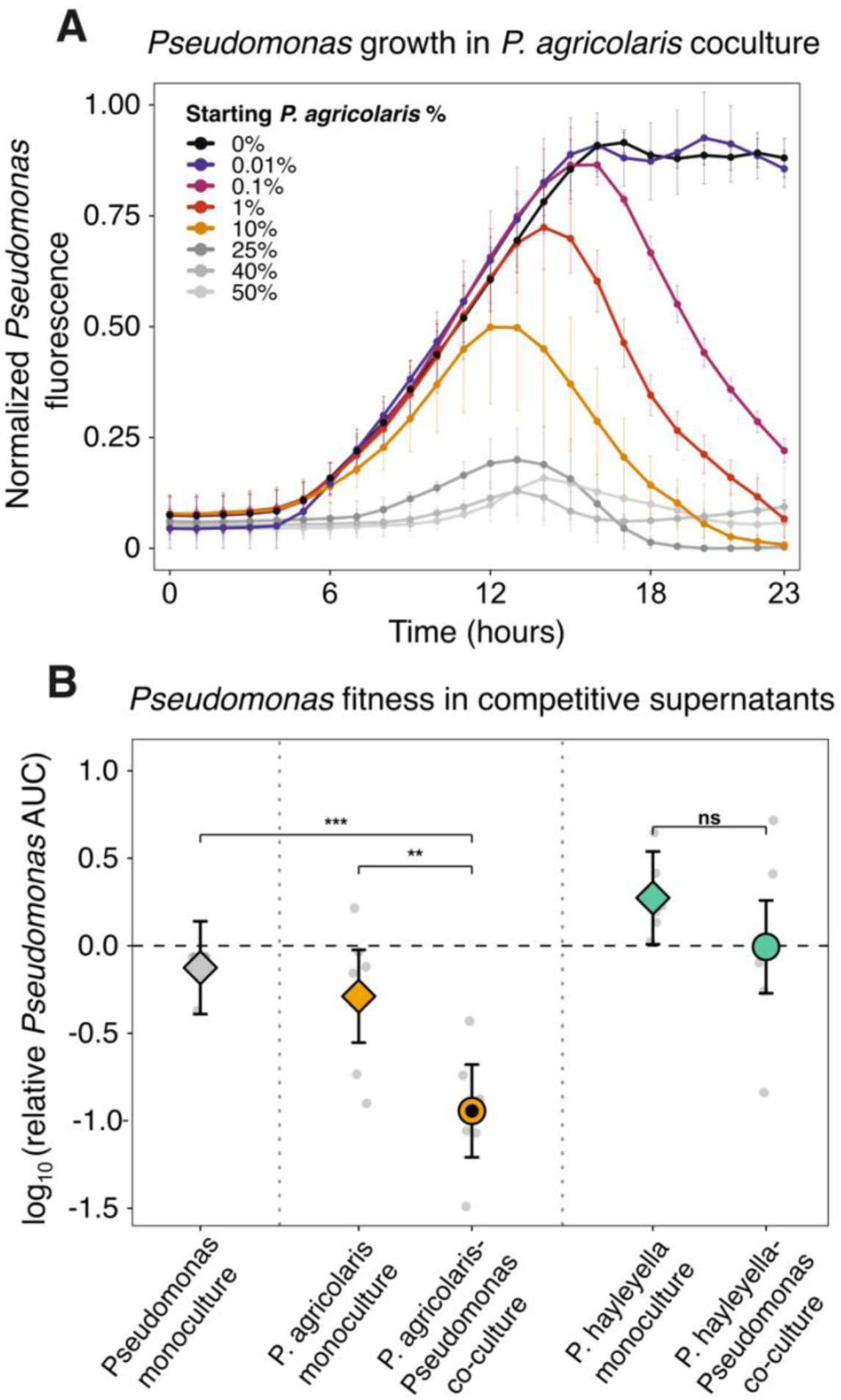
*Pseudomonas* growth is suppressed by *P. agricolaris* in live co-culture and by co-culture-conditioned supernatant. **(A)** Normalised *Pseudomonas* fluorescence during co-culture with varying initial percentages of *P. agricolaris* strain Ba161. Lines show means across biological dates, with error bars indicating SE. Fluorescence was normalised to the maximum corrected fluorescence of same-day *Pseudomonas* monocultures. Suppression occurred even at 0.1% initial Ba161. **(B)** *Pseudomonas* growth in conditioned supernatants from monoculture and co-culture treatments. Coloured symbols show model-estimated means with 95% confidence intervals, while grey points represent date-level observations. Values are log₁₀-transformed AUC relative to *Pseudomonas* monoculture growth in fresh YG, with the dashed line indicating no deviation from this baseline. Black centres within central means indicate significant departures from the baseline. Significance brackets indicate planned contrasts between conditioned-supernatant treatments, with Holm adjustment applied separately across four comparisons against *Pseudomonas* monoculture-conditioned supernatant and two within-species comparisons (** p < 0.01; *** p < 0.001; ns, not significant).

We then tested whether this suppression could be reproduced in the absence of live Ba161 cells using filtered culture supernatants. We grew *Pseudomonas* in the supernatants produced from Ba161, *P. hayleyella* strain Bh11, and *Pseudomonas* monocultures, as well as from Ba161–*Pseudomonas* and Bh11–*Pseudomonas* co-cultures. Both supernatant source and experimental date significantly affected *Pseudomonas* growth (treatment: F₄,₂₀ = 12.81, P < 0.001; date: F₅,₂₀ = 2.94, p = 0.038; Table S7). Planned comparisons revealed that supernatant from the Ba161– *Pseudomonas* co-culture significantly suppressed subsequent *Pseudomonas* growth relative to the *Pseudomonas* monoculture-supernatant control (estimate = −0.818, Holm-adjusted p < 0.001) and Ba161 monoculture supernatant (estimate = −0.655, Holm-adjusted p = 0.003; Table S8). No comparable inhibition was observed with supernatant from Bh11–*Pseudomonas* co-cultures (Fig. 5B; Table S8).

## Discussion

Ecological specialisation can reduce performance outside a favoured environment, either through direct evolutionary trade-offs or through relaxed selection on traits important elsewhere [47–50]. Host adaptation during facultative endosymbiosis presents a striking example, as traits that experience relaxed selection within the host may remain important for survival and competition outside of it [21,51]. Consistent with this possibility, neither streamlined-genome symbiont (*P. bonniea* or *P. hayleyella*) outcompeted any of the tested competitors, and both exhibited lower estimated competitive performance than the non-reduced symbiont *P. auricularis* across all six, with most differences statistically significant (Fig. 2). Contrary to our prediction that amoeba association would generally reduce competitiveness, *P. agricolaris* appeared unusually competitive even among non-amoeba-associated *Paraburkholderia.* Nevertheless, pooled contrasts showed that reduced-genome species performed worse on average than the non-reduced species even when *P. agricolaris* was excluded (Fig. 3). Nevertheless, these differences depended on competitor identity: both reduced species outperformed the non-amoeba-associated group against *Serratia*, despite all losing to *Serratia* (Fig. 2).

Growth-trait analyses helped distinguish changes in the symbionts’ own performance from their effects on competitors. Both reduced-genome species showed greater declines in yield and early maximum growth rate than *P. agricolaris*, although only *P. bonniea* differed from the non-amoeba-associated group in both traits. Conversely, competitors exhibited lower WT-normalised fluorescence yields when paired with *P. agricolaris* than with any other *Paraburkholderia* group (Fig. 4). Together, these findings suggest that *P. agricolaris* combines a greater ability to maintain its own growth with a stronger relative limitation of competitor yield. One explanation for the reduced species’ poorer competitive performance could be their previously documented reduction in carbohydrate metabolic capacity [27]. However, both species grew comparably to *P. agricolaris* in monoculture on glucose-rich medium (Fig. S3), suggesting that poor intrinsic growth under these conditions is insufficient to explain their competitive disadvantage.

An alternative explanation is that reduced competitiveness reflects a trade-off between resistance to amoeba predation and performance during interbacterial competition, in line with a general trade-off between competitive fitness and predation resistance often proposed in bacteria [52–54]. This possibility is especially relevant here because our competitor panel is made up of known high-value prey items for *D. discoideum*, whereas the focal *Paraburkholderia* symbionts range from poor to inedible prey [28,40]. However, the strong competitive performance *of P. agricolaris*, which is also poorly edible, argues against predation resistance alone explaining the competitive disadvantage of the reduced species. Instead, the reduced competitive ability of *P. bonniea* and *P. hayleyella* may reflect ecological costs associated with genome streamlining and relaxed selection for extracellular competitiveness.

Competitive outcomes may depend not only on intrinsic growth characteristics, but also on how organisms respond to and modify their environment during interactions with other species. Experimental studies using *Streptomyces* have demonstrated that encounters with competing bacteria can induce or suppress antagonistic activity depending on competitor identity [55,56]. Complementary work in *Burkholderia thailandensis* has linked interspecific encounters to changes in the production of extracellular metabolites involved in competition [57]. Our finding that Ba161– *Pseudomonas* co-culture generated a more suppressive supernatant than either monoculture suggests that the interaction produced conditions particularly unfavourable for subsequent *Pseudomonas* growth (Fig. 5B). These findings suggest that differences in how *P. agricolaris* and *P. hayleyella* modify their shared environment during competition may contribute to their broader disparity in competitive performance.

Other forms of interbacterial antagonism could also contribute to these competitive outcomes. All three amoeba-associated *Paraburkholderia* are known to possess multiple secretion system complexes with potential roles in interbacterial antagonism [5,58]. This includes a linked type III-type VI secretion system complex uniquely shared across all three species [30]. Both components of this system are upregulated during host infection [37], and the type VI system has been shown to contribute to the horizontal transmission of *P. bonniea* between hosts [34]. This does not rule out an antibacterial function, as a type VI secretion system in *Vibrio cholerae* targets both *Dictyostelium* and other bacteria [59]. Whether the retained *Paraburkholderia* secretion systems contribute to interspecific competition remains unknown. Nevertheless, their retention suggests that the reduced competitiveness of *P. bonniea* and *P. hayleyella* cannot simply be explained by the complete absence of potentially antagonistic machinery.

*D. discoideum* has been found to predate a majority of the culturable prey it co-occurs with [60–62]. A consequence may be that facultative endosymbionts able to resist amoeba predation may not only benefit from physical protection and resource access within the host cell, but also ecological protection from their host as it consumes would-be competitors. Previous work found that *D. discoideum* increased the fitness of *P. hayleyella* during competition with the highly edible bacterium *Klebsiella pneumoniae* in soil mesocosms, but *P. agricolaris* experienced no benefit [32]. Similarly, the six species of competitor bacteria employed here were selected for their established high edibility as *D. discoideum* prey [40], and were isolated from *D. discoideum* fruiting bodies from a location known to contain all three focal *Paraburkholderia* endosymbionts [28,60]. These findings raise the possibility that host predation may reduce interspecies competitive pressure for the reduced-genome symbionts, although the previous mesocosm experiments could not distinguish competitor removal from benefits associated with intracellular residence.

A comparable ecological benefit has been proposed in the *Paramecium bursaria*–*Chlorella* symbiosis. In experimental communities, endosymbiotic *Chlorella* can grow while their ciliate host consumes *Chlamydomonas reinhardtii*, an alga that suppresses the growth of free-living *Chlorella* [63]. Beyond endosymbiosis, mutualistic ants have been shown to prey on aphids that would otherwise outcompete the aphid species they protect, reversing the competitive hierarchy in favour of their mutualistic partners [64]. Together, these examples suggest that consumption of competitors by a host or mutualistic partner can reshape competitive outcomes, potentially altering the selective pressures experienced by symbionts.

Host-mediated protection from competitors could facilitate a form of compensated trait loss, in which an ecological partner supplies a function that would otherwise require an organism to maintain its own capabilities [65]. If *D. discoideum* reduces competitive pressures by consuming other bacteria, selection for its symbionts to maintain their own competitive capabilities may weaken. Such protection could help explain why our reduced-genome species, despite requiring extracellular stages for horizontal transmission, have never been isolated independently from their hosts: their reduced competitive capabilities may make host association increasingly advantageous. However, reduced competitive ability should not be considered an inevitable consequence of adaptation to symbiosis. Even derived bacterial symbionts may retain specialised antagonistic capabilities when these remain advantageous under their particular ecological conditions [66,67].

The evolutionary costs of specialisation may therefore be expressed not only in the environments where organisms can no longer grow, but also in the ecological interactions they are less able to navigate independently. Future work should test whether host-mediated protection relaxes selection on competitive capabilities and establish whether the association between genome streamlining and reduced competitiveness recurs beyond the shared evolutionary lineage of *P. bonniea* and *P. hayleyella*. Such costs could create a form of cryptic dependence, in which symbionts retain the physiological capacity for independent growth while becoming less able to persist in competitive microbial communities. Genome streamlining may therefore reinforce host association before obligate dependence evolves, not because these bacteria cannot grow without their hosts, but because they are less able to compete without them.

## Acknowledgements

We thank Emily Oxender for aiding in early test runs of competition assays, and both Emily and Kaitlyn Torack for help with laboratory work. We thank current and former members of the Strassmann-Queller lab for thoughtful discussions and help with analyses, with specific thanks to both Israt Jahan and P. M. Shreenidhi. Generative AI was used to assist in writing R code for data visualisation and statistical analysis.

## Author Contributions

C.J.S., D.C.Q., and J.E.S. conceived the study, designed experiments, and wrote the manuscript. C.J.S. isolated and transformed bacterial strains, performed the competition assays and the statistical analysis. E.B. transformed bacterial strains and performed the starting-ratio and supernatant assays. All authors approved the final manuscript.

## Data Avaliability

The datasets generated during experimental work here and the R scripts used for data processing, statistical analyses, and figure generation are available on GitHub (https://github.com/mutualistics/paraburkholderia_competition).

