## Supplemental Tables for "*Paraburkholderia* endosymbionts with streamlined genomes are disadvantaged during interspecific competition"

### Supplementary statistical tables

**Table S1: Pairwise contrasts of relative competitive performance within competitor species.** Estimates represent Group 1 − Group 2 differences in $\log_{10}$(monoculture-normalised RFP AUC / GFP AUC), with positive values favouring Group 1. The non-amoeba-associated group (Non-AA) represents the equally weighted mean of *Paraburkholderia fungorum*, *P. unamae* and *P. silvatlantica*. P-values were Holm-adjusted across six contrasts per competitor. Confidence intervals are unadjusted 95% Wald intervals.

| Competitor | Group 1 | Group 2 | $\Delta\log_{10}$ | SE | Lower 95% CI | Upper 95% CI | z | P | P (Holm) |
|---|---|---|---|---|---|---|---|---|---|
| Agrobacterium | Ba | Non-AA | 0.209 | 0.096 | 0.021 | 0.396 | 2.18 | 0.029 | 0.058 |
| Agrobacterium | Ba | Bb | 0.611 | 0.098 | 0.419 | 0.802 | 6.26 | <0.001 | <0.001 |
| Agrobacterium | Ba | Bh | 0.247 | 0.082 | 0.086 | 0.409 | 3.00 | 0.003 | 0.008 |
| Agrobacterium | Bb | Non-AA | -0.402 | 0.085 | -0.568 | -0.236 | -4.75 | <0.001 | <0.001 |
| Agrobacterium | Bb | Bh | -0.364 | 0.071 | -0.503 | -0.224 | -5.11 | <0.001 | <0.001 |
| Agrobacterium | Bh | Non-AA | -0.038 | 0.067 | -0.170 | 0.094 | -0.57 | 0.569 | 0.569 |
| Brucella | Ba | Non-AA | 0.100 | 0.076 | -0.049 | 0.248 | 1.31 | 0.190 | 0.380 |
| Brucella | Ba | Bb | 0.306 | 0.076 | 0.157 | 0.456 | 4.01 | <0.001 | <0.001 |
| Brucella | Ba | Bh | 0.317 | 0.064 | 0.192 | 0.442 | 4.96 | <0.001 | <0.001 |
| Brucella | Bb | Non-AA | -0.207 | 0.066 | -0.336 | -0.078 | -3.14 | 0.002 | 0.005 |
| Brucella | Bb | Bh | 0.010 | 0.055 | -0.097 | 0.117 | 0.19 | 0.852 | 0.852 |
| Brucella | Bh | Non-AA | -0.217 | 0.054 | -0.323 | -0.112 | -4.04 | <0.001 | <0.001 |
| Comamonas | Ba | Non-AA | 0.354 | 0.100 | 0.159 | 0.550 | 3.55 | <0.001 | 0.002 |
| Comamonas | Ba | Bb | 0.256 | 0.101 | 0.057 | 0.455 | 2.53 | 0.012 | 0.046 |
| Comamonas | Ba | Bh | 0.177 | 0.087 | 0.007 | 0.348 | 2.04 | 0.042 | 0.126 |
| Comamonas | Bb | Non-AA | 0.098 | 0.083 | -0.064 | 0.260 | 1.19 | 0.236 | 0.472 |
| Comamonas | Bb | Bh | -0.079 | 0.068 | -0.212 | 0.054 | -1.16 | 0.245 | 0.472 |
| Comamonas | Bh | Non-AA | 0.177 | 0.066 | 0.048 | 0.306 | 2.69 | 0.007 | 0.036 |
| Pseudomonas | Ba | Non-AA | 0.513 | 0.102 | 0.314 | 0.713 | 5.04 | <0.001 | <0.001 |
| Pseudomonas | Ba | Bb | 0.949 | 0.106 | 0.742 | 1.156 | 8.97 | <0.001 | <0.001 |
| Pseudomonas | Ba | Bh | 0.849 | 0.087 | 0.678 | 1.020 | 9.73 | <0.001 | <0.001 |
| Pseudomonas | Bb | Non-AA | -0.436 | 0.089 | -0.610 | -0.261 | -4.90 | <0.001 | <0.001 |
| Pseudomonas | Bb | Bh | -0.100 | 0.075 | -0.247 | 0.047 | -1.34 | 0.181 | 0.181 |
| Pseudomonas | Bh | Non-AA | -0.335 | 0.069 | -0.471 | -0.200 | -4.85 | <0.001 | <0.001 |
| Serratia | Ba | Non-AA | 0.369 | 0.052 | 0.266 | 0.471 | 7.07 | <0.001 | <0.001 |
| Serratia | Ba | Bb | 0.250 | 0.051 | 0.151 | 0.349 | 4.94 | <0.001 | <0.001 |
| Serratia | Ba | Bh | 0.175 | 0.039 | 0.098 | 0.252 | 4.46 | <0.001 | <0.001 |
| Serratia | Bb | Non-AA | 0.119 | 0.048 | 0.024 | 0.213 | 2.46 | 0.014 | 0.028 |
| Serratia | Bb | Bh | -0.075 | 0.038 | -0.150 | 0.001 | -1.95 | 0.052 | 0.052 |
| Serratia | Bh | Non-AA | 0.193 | 0.041 | 0.113 | 0.274 | 4.70 | <0.001 | <0.001 |
| Staphylococcus | Ba | Non-AA | 0.088 | 0.054 | -0.018 | 0.195 | 1.62 | 0.105 | 0.379 |
| Staphylococcus | Ba | Bb | 0.104 | 0.054 | -0.001 | 0.210 | 1.94 | 0.053 | 0.263 |
| Staphylococcus | Ba | Bh | 0.015 | 0.041 | -0.064 | 0.095 | 0.38 | 0.703 | 1.000 |
| Staphylococcus | Bb | Non-AA | -0.016 | 0.052 | -0.118 | 0.086 | -0.31 | 0.756 | 1.000 |

| Competitor | Group 1 | Group 2 | $\Delta log_{10}$ | SE | Lower 95% CI | Upper 95% CI | z | P | P (Holm) |
|---|---|---|---|---|---|---|---|---|---|
| Staphylococcus | Bb | Bh | -0.089 | 0.043 | -0.173 | -0.005 | -2.08 | 0.037 | 0.225 |
| Staphylococcus | Bh | Non-AA | 0.073 | 0.044 | -0.013 | 0.158 | 1.67 | 0.095 | 0.379 |

**Table S2: Model-estimated relative competitive performance tested against parity.** Zero indicates equal monoculture-normalised AUC for the two bacterial partners. The non-amoeba-associated group represents the equally weighted mean of *P. fungorum*, *P. unamae* and *P. silvatlantica*. P-values were Holm-adjusted across 24 tests comprising four *Paraburkholderia* groups and six competitors. Confidence intervals are unadjusted 95% Wald intervals.

| Competitor | Group | Estimate ($log_{10}$) | SE | Lower 95% CI | Upper 95% CI | z | P | P (Holm) |
|---|---|---|---|---|---|---|---|---|
| Agrobacterium | Non-AA | 0.051 | 0.057 | -0.060 | 0.163 | 0.91 | 0.365 | 0.950 |
| Agrobacterium | Ba | 0.260 | 0.077 | 0.109 | 0.411 | 3.38 | <0.001 | 0.007 |
| Agrobacterium | Bb | -0.351 | 0.063 | -0.474 | -0.228 | -5.59 | <0.001 | <0.001 |
| Agrobacterium | Bh | 0.013 | 0.036 | -0.058 | 0.084 | 0.36 | 0.718 | 0.950 |
| Brucella | Non-AA | 0.084 | 0.045 | -0.005 | 0.173 | 1.85 | 0.064 | 0.319 |
| Brucella | Ba | 0.184 | 0.061 | 0.064 | 0.303 | 3.02 | 0.003 | 0.023 |
| Brucella | Bb | -0.123 | 0.048 | -0.216 | -0.029 | -2.56 | 0.010 | 0.062 |
| Brucella | Bh | -0.133 | 0.029 | -0.189 | -0.076 | -4.61 | <0.001 | <0.001 |
| Comamonas | Non-AA | -0.272 | 0.056 | -0.382 | -0.161 | -4.83 | <0.001 | <0.001 |
| Comamonas | Ba | 0.083 | 0.082 | -0.079 | 0.244 | 1.00 | 0.317 | 0.950 |
| Comamonas | Bb | -0.174 | 0.060 | -0.292 | -0.055 | -2.87 | 0.004 | 0.033 |
| Comamonas | Bh | -0.095 | 0.034 | -0.161 | -0.028 | -2.79 | 0.005 | 0.037 |
| Pseudomonas | Non-AA | -0.075 | 0.059 | -0.191 | 0.041 | -1.26 | 0.206 | 0.825 |
| Pseudomonas | Ba | 0.439 | 0.083 | 0.276 | 0.601 | 5.29 | <0.001 | <0.001 |
| Pseudomonas | Bb | -0.510 | 0.066 | -0.641 | -0.380 | -7.69 | <0.001 | <0.001 |
| Pseudomonas | Bh | -0.410 | 0.036 | -0.481 | -0.340 | -11.44 | <0.001 | <0.001 |
| Serratia | Non-AA | -0.590 | 0.035 | -0.658 | -0.522 | -16.97 | <0.001 | <0.001 |
| Serratia | Ba | -0.221 | 0.039 | -0.297 | -0.145 | -5.70 | <0.001 | <0.001 |
| Serratia | Bb | -0.471 | 0.033 | -0.537 | -0.406 | -14.09 | <0.001 | <0.001 |
| Serratia | Bh | -0.396 | 0.022 | -0.439 | -0.353 | -17.94 | <0.001 | <0.001 |
| Staphylococcus | Non-AA | -0.544 | 0.037 | -0.616 | -0.473 | -14.84 | <0.001 | <0.001 |
| Staphylococcus | Ba | -0.456 | 0.040 | -0.535 | -0.378 | -11.37 | <0.001 | <0.001 |
| Staphylococcus | Bb | -0.561 | 0.037 | -0.632 | -0.489 | -15.28 | <0.001 | <0.001 |
| Staphylococcus | Bh | -0.472 | 0.024 | -0.518 | -0.425 | -20.04 | <0.001 | <0.001 |

**Table S3: Comparison of residual-variance structures for the competitive-performance model.** Candidate models shared *Paraburkholderia* species × competitor fixed effects and random intercepts for experimental date and strain but differed in residual-variance structure. Models were compared using maximum likelihood, with ΔAIC and ΔBIC calculated relative to their respective minima. The selected model (bold) was subsequently refitted using REML.

| Residual variance | k | AIC | ΔAIC | BIC | ΔBIC | logLik | Stable | Selected |
|---|---|---|---|---|---|---|---|---|
| **Additive species + competitor** | **49** | **-242.07** | **0.00** | **-63.62** | **0.00** | **170.04** | **Yes** | **Yes** |
| By Paraburkholderia species | 44 | -210.69 | 31.38 | -50.45 | 13.17 | 149.35 | Yes | |
| By competitor | 44 | -174.18 | 67.90 | -13.93 | 49.69 | 131.09 | Yes | |
| Constant | 39 | -143.50 | 98.58 | -1.46 | 62.16 | 110.75 | Yes | |

**Table S4: Pairwise contrasts among *Paraburkholderia* groups for fluorescence-derived yield (*K*) and maximum growth rate ($\mu_{max}$).** Estimates represent Group 1 − Group 2 differences in WT-benchmarked $\log_2$ fold-change, separately for *Paraburkholderia* and competitor responses, averaged equally across six competitors. The non-amoeba-associated group represents the equally weighted mean of *P. fungorum*, *P. unamae* and *P. silvatlantica*. P-values were Holm-adjusted across six contrasts within each response × trait.

| Response | Trait | Group 1 | Group 2 | Δ$\log_2$ FC | SE | Lower 95% CI | Upper 95% CI | z | P | P (Holm) |
|---|---|---|---|---|---|---|---|---|---|---|
| Paraburkholderia | K | Non-AA | Ba | -0.437 | 0.212 | -0.852 | -0.023 | -2.07 | 0.039 | 0.116 |
| Paraburkholderia | K | Non-AA | Bb | 0.664 | 0.214 | 0.245 | 1.084 | 3.10 | 0.002 | 0.008 |
| Paraburkholderia | K | Non-AA | Bh | 0.340 | 0.208 | -0.067 | 0.747 | 1.64 | 0.101 | 0.154 |
| Paraburkholderia | K | Ba | Bb | 1.101 | 0.188 | 0.733 | 1.470 | 5.85 | <0.001 | <0.001 |
| Paraburkholderia | K | Ba | Bh | 0.778 | 0.176 | 0.432 | 1.123 | 4.41 | <0.001 | <0.001 |
| Paraburkholderia | K | Bb | Bh | -0.324 | 0.183 | -0.683 | 0.035 | -1.77 | 0.077 | 0.154 |
| Paraburkholderia | μmax | Non-AA | Ba | -0.067 | 0.165 | -0.390 | 0.255 | -0.41 | 0.682 | 0.682 |
| Paraburkholderia | μmax | Non-AA | Bb | 0.814 | 0.176 | 0.469 | 1.159 | 4.63 | <0.001 | <0.001 |
| Paraburkholderia | μmax | Non-AA | Bh | 0.320 | 0.165 | -0.003 | 0.644 | 1.94 | 0.052 | 0.104 |
| Paraburkholderia | μmax | Ba | Bb | 0.882 | 0.159 | 0.570 | 1.194 | 5.54 | <0.001 | <0.001 |
| Paraburkholderia | μmax | Ba | Bh | 0.388 | 0.144 | 0.106 | 0.670 | 2.69 | 0.007 | 0.021 |
| Paraburkholderia | μmax | Bb | Bh | -0.494 | 0.160 | -0.807 | -0.181 | -3.10 | 0.002 | 0.008 |
| Competitor | K | Non-AA | Ba | 0.542 | 0.087 | 0.371 | 0.713 | 6.21 | <0.001 | <0.001 |
| Competitor | K | Non-AA | Bb | 0.062 | 0.079 | -0.094 | 0.217 | 0.78 | 0.437 | 0.874 |
| Competitor | K | Non-AA | Bh | 0.085 | 0.077 | -0.066 | 0.236 | 1.10 | 0.270 | 0.811 |
| Competitor | K | Ba | Bb | -0.480 | 0.076 | -0.629 | -0.332 | -6.34 | <0.001 | <0.001 |
| Competitor | K | Ba | Bh | -0.457 | 0.068 | -0.590 | -0.324 | -6.76 | <0.001 | <0.001 |
| Competitor | K | Bb | Bh | 0.023 | 0.064 | -0.102 | 0.149 | 0.37 | 0.714 | 0.874 |
| Competitor | μmax | Non-AA | Ba | 0.103 | 0.076 | -0.047 | 0.253 | 1.34 | 0.180 | 0.359 |
| Competitor | μmax | Non-AA | Bb | -0.226 | 0.085 | -0.392 | -0.060 | -2.66 | 0.008 | 0.039 |
| Competitor | μmax | Non-AA | Bh | -0.054 | 0.076 | -0.203 | 0.095 | -0.71 | 0.477 | 0.477 |
| Competitor | μmax | Ba | Bb | -0.329 | 0.078 | -0.482 | -0.175 | -4.20 | <0.001 | <0.001 |
| Competitor | μmax | Ba | Bh | -0.157 | 0.068 | -0.289 | -0.024 | -2.32 | 0.020 | 0.081 |
| Competitor | μmax | Bb | Bh | 0.172 | 0.078 | 0.019 | 0.325 | 2.21 | 0.027 | 0.082 |

**Table S5: Model-estimated fluorescence-derived growth traits relative to wild-type controls.** Estimates represent WT-benchmarked $log_2$ fold-changes in *K* and μmax for both partners, averaged equally across six competitors. Zero indicates no deviation from the matched labelled–unlabelled wild-type (WT) co-culture baseline. P-values were Holm-adjusted across four *Paraburkholderia* groups within each response × trait.

| Response | Trait | Group | Estimate ($log_2$FC) | SE | Lower 95% CI | Upper 95% CI | z | P | P (Holm) |
|---|---|---|---|---|---|---|---|---|---|
| Paraburkholderia | K | Non-AA | -0.355 | 0.166 | -0.679 | -0.030 | -2.14 | 0.032 | 0.064 |
| Paraburkholderia | K | Ba | 0.082 | 0.132 | -0.176 | 0.340 | 0.62 | 0.532 | 0.532 |
| Paraburkholderia | K | Bb | -1.019 | 0.136 | -1.285 | -0.753 | -7.52 | <0.001 | <0.001 |
| Paraburkholderia | K | Bh | -0.695 | 0.125 | -0.941 | -0.449 | -5.54 | <0.001 | <0.001 |
| Paraburkholderia | μmax | Non-AA | -0.061 | 0.127 | -0.311 | 0.189 | -0.48 | 0.633 | 1.000 |
| Paraburkholderia | μmax | Ba | 0.007 | 0.104 | -0.198 | 0.211 | 0.06 | 0.950 | 1.000 |
| Paraburkholderia | μmax | Bb | -0.875 | 0.122 | -1.113 | -0.637 | -7.20 | <0.001 | <0.001 |
| Paraburkholderia | μmax | Bh | -0.381 | 0.105 | -0.586 | -0.176 | -3.64 | <0.001 | <0.001 |
| Competitor | K | Non-AA | 0.574 | 0.063 | 0.451 | 0.697 | 9.16 | <0.001 | <0.001 |
| Competitor | K | Ba | 0.032 | 0.061 | -0.087 | 0.151 | 0.52 | 0.601 | 0.601 |
| Competitor | K | Bb | 0.512 | 0.049 | 0.417 | 0.607 | 10.54 | <0.001 | <0.001 |
| Competitor | K | Bh | 0.489 | 0.045 | 0.400 | 0.577 | 10.82 | <0.001 | <0.001 |
| Competitor | μmax | Non-AA | 0.225 | 0.059 | 0.110 | 0.340 | 3.84 | <0.001 | <0.001 |
| Competitor | μmax | Ba | 0.123 | 0.049 | 0.027 | 0.219 | 2.50 | 0.012 | 0.012 |
| Competitor | μmax | Bb | 0.451 | 0.061 | 0.331 | 0.571 | 7.36 | <0.001 | <0.001 |
| Competitor | μmax | Bh | 0.279 | 0.048 | 0.185 | 0.374 | 5.79 | <0.001 | <0.001 |

**Table S6: Comparison of mixed-effects model structures for fluorescence-derived yield and maximum growth rate.** Candidate models differed in fixed effects, random intercepts and residual-variance structure, and were compared using maximum likelihood separately for each response × trait. Models are ranked by AIC, with selected models shown in bold and subsequently refitted using REML. ΔAIC and ΔBIC are calculated relative to their respective minima. Model structures are defined below the table.

| Response | Trait | Model | n | k | AIC | ΔAIC | BIC | ΔBIC | logLik | Stable | Selected |
|---|---|---|---|---|---|---|---|---|---|---|---|
| **Paraburkholderia** | **K** | **M6** | **282** | **49** | **352.51** | **0.00** | **530.97** | **0.35** | **-127.26** | **Yes** | **Yes** |
| Paraburkholderia | K | M5 | 282 | 48 | 366.96 | 14.45 | 541.78 | 11.16 | -135.48 | Yes | |
| Paraburkholderia | K | M3 | 282 | 43 | 374.02 | 21.50 | 530.62 | 0.00 | -144.01 | Yes | |
| Paraburkholderia | K | M4 | 282 | 43 | 405.67 | 53.16 | 562.27 | 31.65 | -159.84 | Yes | |
| Paraburkholderia | K | M2 | 282 | 39 | 414.12 | 61.61 | 556.16 | 25.54 | -168.06 | Yes | |
| Paraburkholderia | K | M1 | 282 | 38 | 428.33 | 75.82 | 566.72 | 36.10 | -176.16 | Yes | |
| Paraburkholderia | K | M0 | 282 | 13 | 585.22 | 232.71 | 632.57 | 101.95 | -279.61 | Yes | |
| **Paraburkholderia** | **μmax** | **M6** | **269** | **49** | **269.71** | **0.00** | **445.85** | **0.00** | **-85.85** | **Yes** | **Yes** |
| Paraburkholderia | μmax | M5 | 269 | 48 | 288.06 | 18.36 | 460.61 | 14.76 | -96.03 | Yes | |
| Paraburkholderia | μmax | M4 | 269 | 43 | 297.89 | 28.18 | 452.46 | 6.61 | -105.94 | Yes | |
| Paraburkholderia | μmax | M3 | 269 | 43 | 366.69 | 96.98 | 521.26 | 75.41 | -140.34 | Yes | |
| Paraburkholderia | μmax | M2 | 269 | 39 | 379.61 | 109.90 | 519.80 | 73.96 | -150.80 | Yes | |
| Paraburkholderia | μmax | M1 | 269 | 38 | 379.96 | 110.26 | 516.56 | 70.72 | -151.98 | Yes | |
| Paraburkholderia | μmax | M0 | 269 | 13 | 447.26 | 177.55 | 493.99 | 48.14 | -210.63 | Yes | |
| **Competitor** | **K** | **M5** | **282** | **48** | **196.01** | **0.00** | **370.82** | **68.05** | **-50.00** | **Yes** | **Yes** |
| Competitor | K | M6 | 282 | 49 | 198.01 | 2.00 | 376.46 | 73.69 | -50.00 | Yes | |
| Competitor | K | M4 | 282 | 43 | 216.78 | 20.77 | 373.38 | 70.61 | -65.39 | Yes | |
| Competitor | K | M3 | 282 | 43 | 228.66 | 32.65 | 385.26 | 82.49 | -71.33 | Yes | |
| Competitor | K | M0 | 282 | 13 | 255.43 | 59.42 | 302.77 | 0.00 | -114.71 | Yes | |
| Competitor | K | M1 | 282 | 38 | 261.94 | 65.93 | 400.34 | 97.57 | -92.97 | Yes | |
| Competitor | K | M2 | 282 | 39 | 263.94 | 67.93 | 405.98 | 103.21 | -92.97 | Yes | |
| **Competitor** | **μmax** | **M5** | **282** | **48** | **40.26** | **0.00** | **215.07** | **0.00** | **27.87** | **Yes** | **Yes** |
| Competitor | μmax | M6 | 282 | 49 | 42.26 | 2.00 | 220.72 | 5.64 | 27.87 | Yes | |
| Competitor | μmax | M4 | 282 | 43 | 58.82 | 18.56 | 215.43 | 0.35 | 13.59 | Yes | |
| Competitor | μmax | M3 | 282 | 43 | 317.35 | 277.09 | 473.95 | 258.88 | -115.67 | Yes | |
| Competitor | μmax | M0 | 282 | 13 | 362.70 | 322.44 | 410.04 | 194.97 | -168.35 | Yes | |
| Competitor | μmax | M1 | 282 | 38 | 369.74 | 329.48 | 508.13 | 293.06 | -146.87 | Yes | |
| Competitor | μmax | M2 | 282 | 39 | 371.74 | 331.48 | 513.77 | 298.70 | -146.87 | Yes | |

**Model definitions:** M0 included additive *Paraburkholderia* species and competitor effects, a date random intercept and constant residual variance. M1 included the species × competitor interaction; M2 additionally included a strain random intercept. M3 and M4 allowed residual variance to vary by *Paraburkholderia* species or competitor, respectively. M5 included additive species- and competitor-specific residual dispersion, and M6 additionally included a strain random intercept. Models M1 and M3–M5 included a date random intercept.

**Table S7: Effects of conditioned-supernatant treatment and experimental date on *Pseudomonas* growth.** Model terms were evaluated using single-term deletion F-tests from a date-blocked linear model of $\log_{10}$(monoculture-normalised *Pseudomonas* GFP AUC). Treatment comprised five conditioned-supernatant preparations measured across six experimental dates (30 observations; residual df = 20).

| Term | df (numerator) | df (denominator) | F | P |
|---|---|---|---|---|
| Treatment | 4 | 20 | 12.81 | <0.001 |
| Date | 5 | 20 | 2.94 | 0.038 |

**Table S8: Planned contrasts and treatment means for the conditioned-supernatant experiment.** Estimates represent differences between treatments or individual treatment means of $\log_{10}$-transformed *Pseudomonas* GFP AUC, normalised to *Pseudomonas* monoculture growth in fresh YG medium. Ba denotes *Paraburkholderia agricolaris* strain Ba161; Bh, *Paraburkholderia hayleyella* strain Bh11; and Pk, *Pseudomonas kuykendallii* strain 14P 8.1.1. P-values were Holm-adjusted separately across four comparisons against Pk monoculture-conditioned supernatant, two within-species monoculture–co-culture comparisons, and five treatment means tested against zero. Confidence intervals are unadjusted 95% intervals.

| Family | Comparison | Estimate ($\log_{10}$) | SE | Lower 95% CI | Upper 95% CI | t | P | P (Holm) |
|---|---|---|---|---|---|---|---|---|
| Against Pk monoculture (4) | Ba monoculture − Pk monoculture | -0.163 | 0.180 | -0.538 | 0.212 | -0.91 | 0.375 | 0.750 |
| Against Pk monoculture (4) | Ba–Pk co-culture − Pk monoculture | -0.818 | 0.180 | -1.193 | -0.444 | -4.55 | <0.001 | <0.001 |
| Against Pk monoculture (4) | Bh monoculture − Pk monoculture | 0.399 | 0.180 | 0.024 | 0.774 | 2.22 | 0.038 | 0.114 |
| Against Pk monoculture (4) | Bh–Pk co-culture − Pk monoculture | 0.119 | 0.180 | -0.256 | 0.494 | 0.66 | 0.515 | 0.750 |
| Within symbiont species (2) | Ba–Pk co-culture − Ba monoculture | -0.655 | 0.180 | -1.030 | -0.280 | -3.65 | 0.002 | 0.003 |
| Within symbiont species (2) | Bh–Pk co-culture − Bh monoculture | -0.280 | 0.180 | -0.655 | 0.095 | -1.56 | 0.135 | 0.135 |
| Treatment means against zero (5) | Pk mono | -0.125 | 0.127 | -0.390 | 0.140 | -0.99 | 0.335 | 0.670 |
| Treatment means against zero (5) | Ba mono | -0.289 | 0.127 | -0.554 | -0.024 | -2.27 | 0.034 | 0.137 |
| Treatment means against zero (5) | Ba coculture | -0.944 | 0.127 | -1.209 | -0.679 | -7.43 | <0.001 | <0.001 |
| Treatment means against zero (5) | Bh mono | 0.274 | 0.127 | 0.009 | 0.539 | 2.15 | 0.044 | 0.137 |
| Treatment means against zero (5) | Bh coculture | -0.006 | 0.127 | -0.271 | 0.259 | -0.05 | 0.961 | 0.961 |

**Table S9: Pooled comparisons of relative competitive performance by genome-reduction status.** Contrasts compare four non-reduced species against two reduced species, and three non-amoeba-associated species against the two reduced species. Estimates represent differences in the $\log_{10}$ ratio of monoculture-normalised RFP to GFP AUC, averaged equally across six competitors. Positive values favour non-reduced species. P-values were Holm-adjusted across the two planned contrasts.

| Contrast | $\Delta\log_{10}$ | SE | Lower 95% CI | Upper 95% CI | Fold difference | z | P | P (Holm) |
|---|---|---|---|---|---|---|---|---|
| 4 non-reduced-genome species vs Bb/Bh | 0.151 | 0.030 | 0.091 | 0.210 | 1.41 | 4.99 | <0.001 | <0.001 |
| 3 non-AA species vs Bb/Bh | 0.083 | 0.037 | 0.010 | 0.155 | 1.21 | 2.23 | 0.026 | 0.026 |

**Table S10: Competitor-specific fluorescence-derived growth traits relative to wild-type controls.** Model-estimated WT-benchmarked $\log_2$ fold-changes in *K* and μmax are shown for both partners across six competitors. Zero indicates no deviation from the matched labelled–unlabelled WT co-culture baseline. The non-amoeba-associated group represents the equally weighted mean of *P. fungorum*, *P. unamae* and *P. silvatlantica*. P-values were Holm-adjusted across 24 tests within each response × trait.

| Response | Trait | Competitor | Group | Estimate ($\log_2$FC) | SE | Lower 95% CI | Upper 95% CI | z | P | P (Holm) |
|---|---|---|---|---|---|---|---|---|---|---|
| Paraburkholderia | K | Agrobacterium | Non-AA | 0.632 | 0.260 | 0.123 | 1.140 | 2.43 | 0.015 | 0.135 |
| Paraburkholderia | K | Agrobacterium | Ba | 0.719 | 0.183 | 0.361 | 1.077 | 3.94 | <0.001 | 0.001 |
| Paraburkholderia | K | Agrobacterium | Bb | -1.123 | 0.198 | -1.511 | -0.736 | -5.68 | <0.001 | <0.001 |
| Paraburkholderia | K | Agrobacterium | Bh | -0.027 | 0.150 | -0.321 | 0.268 | -0.18 | 0.860 | 1.000 |
| Paraburkholderia | K | Brucella | Non-AA | 0.624 | 0.186 | 0.258 | 0.989 | 3.35 | <0.001 | 0.010 |
| Paraburkholderia | K | Brucella | Ba | 0.548 | 0.146 | 0.262 | 0.833 | 3.76 | <0.001 | 0.002 |
| Paraburkholderia | K | Brucella | Bb | -0.200 | 0.150 | -0.493 | 0.093 | -1.34 | 0.182 | 1.000 |
| Paraburkholderia | K | Brucella | Bh | -0.344 | 0.131 | -0.601 | -0.087 | -2.62 | 0.009 | 0.089 |
| Paraburkholderia | K | Comamonas | Non-AA | -0.001 | 0.241 | -0.475 | 0.472 | -0.01 | 0.995 | 1.000 |
| Paraburkholderia | K | Comamonas | Ba | 0.475 | 0.179 | 0.124 | 0.827 | 2.65 | 0.008 | 0.089 |
| Paraburkholderia | K | Comamonas | Bb | -0.213 | 0.183 | -0.571 | 0.146 | -1.16 | 0.245 | 1.000 |
| Paraburkholderia | K | Comamonas | Bh | -0.098 | 0.142 | -0.377 | 0.182 | -0.69 | 0.493 | 1.000 |
| Paraburkholderia | K | Pseudomonas | Non-AA | -0.630 | 0.276 | -1.170 | -0.090 | -2.29 | 0.022 | 0.178 |
| Paraburkholderia | K | Pseudomonas | Ba | 0.201 | 0.193 | -0.178 | 0.580 | 1.04 | 0.298 | 1.000 |
| Paraburkholderia | K | Pseudomonas | Bb | -1.423 | 0.209 | -1.833 | -1.013 | -6.81 | <0.001 | <0.001 |
| Paraburkholderia | K | Pseudomonas | Bh | -1.166 | 0.150 | -1.460 | -0.872 | -7.78 | <0.001 | <0.001 |
| Paraburkholderia | K | Serratia | Non-AA | -1.346 | 0.228 | -1.794 | -0.899 | -5.90 | <0.001 | <0.001 |
| Paraburkholderia | K | Serratia | Ba | -0.280 | 0.169 | -0.611 | 0.052 | -1.65 | 0.099 | 0.691 |
| Paraburkholderia | K | Serratia | Bb | -1.261 | 0.178 | -1.610 | -0.913 | -7.09 | <0.001 | <0.001 |
| Paraburkholderia | K | Serratia | Bh | -1.031 | 0.139 | -1.303 | -0.759 | -7.42 | <0.001 | <0.001 |
| Paraburkholderia | K | Staphylococcus | Non-AA | -1.407 | 0.215 | -1.829 | -0.985 | -6.54 | <0.001 | <0.001 |
| Paraburkholderia | K | Staphylococcus | Ba | -1.170 | 0.158 | -1.479 | -0.861 | -7.43 | <0.001 | <0.001 |
| Paraburkholderia | K | Staphylococcus | Bb | -1.895 | 0.170 | -2.229 | -1.562 | -11.15 | <0.001 | <0.001 |
| Paraburkholderia | K | Staphylococcus | Bh | -1.507 | 0.136 | -1.774 | -1.240 | -11.05 | <0.001 | <0.001 |
| Paraburkholderia | μmax | Agrobacterium | Non-AA | 0.255 | 0.134 | -0.007 | 0.517 | 1.91 | 0.056 | 0.617 |
| Paraburkholderia | μmax | Agrobacterium | Ba | 0.409 | 0.111 | 0.192 | 0.626 | 3.70 | <0.001 | 0.004 |
| Paraburkholderia | μmax | Agrobacterium | Bb | -0.159 | 0.130 | -0.413 | 0.095 | -1.23 | 0.220 | 1.000 |
| Paraburkholderia | μmax | Agrobacterium | Bh | 0.185 | 0.114 | -0.040 | 0.409 | 1.61 | 0.106 | 1.000 |
| Paraburkholderia | μmax | Brucella | Non-AA | 0.138 | 0.119 | -0.095 | 0.372 | 1.16 | 0.245 | 1.000 |
| Paraburkholderia | μmax | Brucella | Ba | 0.297 | 0.101 | 0.099 | 0.495 | 2.94 | 0.003 | 0.046 |
| Paraburkholderia | μmax | Brucella | Bb | -0.335 | 0.104 | -0.538 | -0.132 | -3.24 | 0.001 | 0.019 |
| Paraburkholderia | μmax | Brucella | Bh | -0.139 | 0.100 | -0.335 | 0.057 | -1.39 | 0.163 | 1.000 |
| Paraburkholderia | μmax | Comamonas | Non-AA | 0.150 | 0.139 | -0.122 | 0.422 | 1.08 | 0.280 | 1.000 |
| Paraburkholderia | μmax | Comamonas | Ba | 0.065 | 0.116 | -0.163 | 0.293 | 0.56 | 0.577 | 1.000 |
| Paraburkholderia | μmax | Comamonas | Bb | -0.202 | 0.126 | -0.449 | 0.044 | -1.61 | 0.108 | 1.000 |
| Paraburkholderia | μmax | Comamonas | Bh | -0.076 | 0.115 | -0.301 | 0.149 | -0.66 | 0.509 | 1.000 |
| Paraburkholderia | μmax | Pseudomonas | Non-AA | 0.199 | 0.208 | -0.209 | 0.606 | 0.96 | 0.339 | 1.000 |
| Paraburkholderia | μmax | Pseudomonas | Ba | 0.454 | 0.166 | 0.129 | 0.779 | 2.74 | 0.006 | 0.079 |
| Paraburkholderia | μmax | Pseudomonas | Bb | -1.361 | 0.221 | -1.795 | -0.927 | -6.14 | <0.001 | <0.001 |
| Paraburkholderia | μmax | Pseudomonas | Bh | -0.632 | 0.167 | -0.959 | -0.305 | -3.78 | <0.001 | 0.003 |
| Paraburkholderia | μmax | Serratia | Non-AA | -0.590 | 0.195 | -0.971 | -0.208 | -3.03 | 0.002 | 0.037 |

| Response | Trait | Competitor | Group | Estimate ($log_2FC$) | SE | Lower 95% CI | Upper 95% CI | z | P | P (Holm) |
|---|---|---|---|---|---|---|---|---|---|---|
| Paraburkholderia | µmax | Serratia | Ba | -0.226 | 0.160 | -0.538 | 0.087 | -1.41 | 0.158 | 1.000 |
| Paraburkholderia | µmax | Serratia | Bb | -0.911 | 0.198 | -1.299 | -0.524 | -4.61 | <0.001 | <0.001 |
| Paraburkholderia | µmax | Serratia | Bh | -0.657 | 0.158 | -0.966 | -0.348 | -4.17 | <0.001 | <0.001 |
| Paraburkholderia | µmax | Staphylococcus | Non-AA | -0.518 | 0.258 | -1.024 | -0.012 | -2.01 | 0.045 | 0.536 |
| Paraburkholderia | µmax | Staphylococcus | Ba | -0.960 | 0.204 | -1.359 | -0.560 | -4.71 | <0.001 | <0.001 |
| Paraburkholderia | µmax | Staphylococcus | Bb | -2.282 | 0.356 | -2.980 | -1.585 | -6.41 | <0.001 | <0.001 |
| Paraburkholderia | µmax | Staphylococcus | Bh | -0.968 | 0.206 | -1.371 | -0.564 | -4.70 | <0.001 | <0.001 |
| Competitor | K | Agrobacterium | Non-AA | -0.041 | 0.123 | -0.281 | 0.200 | -0.33 | 0.741 | 1.000 |
| Competitor | K | Agrobacterium | Ba | -0.788 | 0.151 | -1.083 | -0.493 | -5.23 | <0.001 | <0.001 |
| Competitor | K | Agrobacterium | Bb | -0.157 | 0.102 | -0.358 | 0.043 | -1.54 | 0.124 | 0.618 |
| Competitor | K | Agrobacterium | Bh | -0.334 | 0.100 | -0.529 | -0.139 | -3.35 | <0.001 | 0.009 |
| Competitor | K | Brucella | Non-AA | 0.098 | 0.098 | -0.094 | 0.290 | 1.00 | 0.318 | 1.000 |
| Competitor | K | Brucella | Ba | -0.346 | 0.123 | -0.588 | -0.104 | -2.80 | 0.005 | 0.045 |
| Competitor | K | Brucella | Bb | 0.262 | 0.081 | 0.103 | 0.421 | 3.24 | 0.001 | 0.012 |
| Competitor | K | Brucella | Bh | 0.203 | 0.077 | 0.051 | 0.355 | 2.62 | 0.009 | 0.070 |
| Competitor | K | Comamonas | Non-AA | 0.587 | 0.114 | 0.364 | 0.811 | 5.15 | <0.001 | <0.001 |
| Competitor | K | Comamonas | Ba | -0.145 | 0.150 | -0.439 | 0.149 | -0.97 | 0.334 | 1.000 |
| Competitor | K | Comamonas | Bb | 0.236 | 0.093 | 0.055 | 0.418 | 2.55 | 0.011 | 0.075 |
| Competitor | K | Comamonas | Bh | 0.215 | 0.088 | 0.042 | 0.389 | 2.44 | 0.015 | 0.088 |
| Competitor | K | Pseudomonas | Non-AA | 0.639 | 0.140 | 0.364 | 0.914 | 4.55 | <0.001 | <0.001 |
| Competitor | K | Pseudomonas | Ba | 0.136 | 0.180 | -0.216 | 0.488 | 0.76 | 0.449 | 1.000 |
| Competitor | K | Pseudomonas | Bb | 0.982 | 0.119 | 0.750 | 1.215 | 8.28 | <0.001 | <0.001 |
| Competitor | K | Pseudomonas | Bh | 0.872 | 0.109 | 0.659 | 1.085 | 8.03 | <0.001 | <0.001 |
| Competitor | K | Serratia | Non-AA | 0.990 | 0.062 | 0.868 | 1.112 | 15.92 | <0.001 | <0.001 |
| Competitor | K | Serratia | Ba | 0.560 | 0.064 | 0.434 | 0.686 | 8.70 | <0.001 | <0.001 |
| Competitor | K | Serratia | Bb | 0.787 | 0.051 | 0.686 | 0.888 | 15.31 | <0.001 | <0.001 |
| Competitor | K | Serratia | Bh | 0.850 | 0.047 | 0.758 | 0.943 | 18.03 | <0.001 | <0.001 |
| Competitor | K | Staphylococcus | Non-AA | 1.169 | 0.085 | 1.003 | 1.335 | 13.81 | <0.001 | <0.001 |
| Competitor | K | Staphylococcus | Ba | 0.774 | 0.095 | 0.587 | 0.961 | 8.11 | <0.001 | <0.001 |
| Competitor | K | Staphylococcus | Bb | 0.963 | 0.072 | 0.822 | 1.103 | 13.45 | <0.001 | <0.001 |
| Competitor | K | Staphylococcus | Bh | 1.125 | 0.064 | 0.999 | 1.251 | 17.48 | <0.001 | <0.001 |
| Competitor | µmax | Agrobacterium | Non-AA | -0.111 | 0.185 | -0.474 | 0.252 | -0.60 | 0.550 | 1.000 |
| Competitor | µmax | Agrobacterium | Ba | -0.377 | 0.150 | -0.671 | -0.083 | -2.51 | 0.012 | 0.107 |
| Competitor | µmax | Agrobacterium | Bb | 0.544 | 0.203 | 0.146 | 0.942 | 2.68 | 0.007 | 0.073 |
| Competitor | µmax | Agrobacterium | Bh | -0.010 | 0.158 | -0.320 | 0.300 | -0.06 | 0.951 | 1.000 |
| Competitor | µmax | Brucella | Non-AA | 0.239 | 0.063 | 0.116 | 0.362 | 3.82 | <0.001 | 0.001 |
| Competitor | µmax | Brucella | Ba | -0.051 | 0.054 | -0.157 | 0.055 | -0.94 | 0.348 | 1.000 |
| Competitor | µmax | Brucella | Bb | 0.437 | 0.067 | 0.304 | 0.569 | 6.47 | <0.001 | <0.001 |
| Competitor | µmax | Brucella | Bh | 0.525 | 0.055 | 0.417 | 0.633 | 9.55 | <0.001 | <0.001 |
| Competitor | µmax | Comamonas | Non-AA | 0.200 | 0.263 | -0.316 | 0.717 | 0.76 | 0.447 | 1.000 |
| Competitor | µmax | Comamonas | Ba | 0.379 | 0.230 | -0.072 | 0.830 | 1.65 | 0.099 | 0.696 |
| Competitor | µmax | Comamonas | Bb | 0.603 | 0.277 | 0.059 | 1.147 | 2.17 | 0.030 | 0.237 |
| Competitor | µmax | Comamonas | Bh | -0.054 | 0.216 | -0.478 | 0.370 | -0.25 | 0.802 | 1.000 |
| Competitor | µmax | Pseudomonas | Non-AA | 0.530 | 0.058 | 0.415 | 0.644 | 9.10 | <0.001 | <0.001 |
| Competitor | µmax | Pseudomonas | Ba | 0.654 | 0.048 | 0.559 | 0.749 | 13.54 | <0.001 | <0.001 |
| Competitor | µmax | Pseudomonas | Bb | 0.626 | 0.062 | 0.503 | 0.748 | 10.01 | <0.001 | <0.001 |
| Competitor | µmax | Pseudomonas | Bh | 0.717 | 0.049 | 0.621 | 0.813 | 14.59 | <0.001 | <0.001 |
| Competitor | µmax | Serratia | Non-AA | 0.293 | 0.039 | 0.217 | 0.369 | 7.52 | <0.001 | <0.001 |

| Response | Trait | Competitor | Group | Estimate ($log_2FC$) | SE | Lower 95% CI | Upper 95% CI | z | P | P (Holm) |
|---|---|---|---|---|---|---|---|---|---|---|
| Competitor | µmax | Serratia | Ba | -0.043 | 0.034 | -0.109 | 0.023 | -1.29 | 0.198 | 1.000 |
| Competitor | µmax | Serratia | Bb | 0.169 | 0.041 | 0.089 | 0.248 | 4.14 | <0.001 | <0.001 |
| Competitor | µmax | Serratia | Bh | 0.161 | 0.033 | 0.097 | 0.225 | 4.94 | <0.001 | <0.001 |
| Competitor | µmax | Staphylococcus | Non-AA | 0.201 | 0.031 | 0.141 | 0.261 | 6.55 | <0.001 | <0.001 |
| Competitor | µmax | Staphylococcus | Ba | 0.174 | 0.024 | 0.127 | 0.221 | 7.23 | <0.001 | <0.001 |
| Competitor | µmax | Staphylococcus | Bb | 0.330 | 0.031 | 0.268 | 0.391 | 10.47 | <0.001 | <0.001 |
| Competitor | µmax | Staphylococcus | Bh | 0.336 | 0.025 | 0.287 | 0.386 | 13.33 | <0.001 | <0.001 |

**Table S11: Competitor-specific pairwise contrasts in fluorescence-derived growth traits.** Estimates represent Group 1 − Group 2 differences in WT-benchmarked $\log_2$ fold-change for *K* and μmax, separately for each competitor and both fluorescent partners. Positive values favour Group 1. The non-amoeba-associated group represents the equally weighted mean of *P. fungorum*, *P. unamae* and *P. silvatlantica*. P-values were Holm-adjusted across six contrasts within each response × trait × competitor.

| Response | Trait | Competitor | Group 1 | Group 2 | $\Delta\log_2$FC | SE | Lower 95% CI | Upper 95% CI | z | P | P (Holm) |
|---|---|---|---|---|---|---|---|---|---|---|---|
| Paraburkholderia | K | Agrobacterium | Non-AA | Ba | -0.087 | 0.317 | -0.709 | 0.535 | -0.28 | 0.783 | 0.783 |
| Paraburkholderia | K | Agrobacterium | Non-AA | Bb | 1.755 | 0.326 | 1.115 | 2.394 | 5.38 | <0.001 | <0.001 |
| Paraburkholderia | K | Agrobacterium | Non-AA | Bh | 0.658 | 0.300 | 0.070 | 1.246 | 2.19 | 0.028 | 0.056 |
| Paraburkholderia | K | Agrobacterium | Ba | Bb | 1.842 | 0.266 | 1.321 | 2.364 | 6.92 | <0.001 | <0.001 |
| Paraburkholderia | K | Agrobacterium | Ba | Bh | 0.746 | 0.230 | 0.294 | 1.197 | 3.24 | 0.001 | 0.004 |
| Paraburkholderia | K | Agrobacterium | Bb | Bh | -1.097 | 0.246 | -1.579 | -0.614 | -4.46 | <0.001 | <0.001 |
| Paraburkholderia | K | Brucella | Non-AA | Ba | 0.076 | 0.237 | -0.388 | 0.540 | 0.32 | 0.748 | 0.933 |
| Paraburkholderia | K | Brucella | Non-AA | Bb | 0.824 | 0.239 | 0.355 | 1.292 | 3.44 | <0.001 | 0.002 |
| Paraburkholderia | K | Brucella | Non-AA | Bh | 0.967 | 0.228 | 0.521 | 1.414 | 4.24 | <0.001 | <0.001 |
| Paraburkholderia | K | Brucella | Ba | Bb | 0.747 | 0.208 | 0.340 | 1.154 | 3.60 | <0.001 | 0.001 |
| Paraburkholderia | K | Brucella | Ba | Bh | 0.891 | 0.189 | 0.520 | 1.262 | 4.71 | <0.001 | <0.001 |
| Paraburkholderia | K | Brucella | Bb | Bh | 0.144 | 0.198 | -0.243 | 0.531 | 0.73 | 0.466 | 0.933 |
| Paraburkholderia | K | Comamonas | Non-AA | Ba | -0.477 | 0.301 | -1.066 | 0.113 | -1.58 | 0.113 | 0.452 |
| Paraburkholderia | K | Comamonas | Non-AA | Bb | 0.211 | 0.303 | -0.383 | 0.805 | 0.70 | 0.486 | 1.000 |
| Paraburkholderia | K | Comamonas | Non-AA | Bh | 0.096 | 0.280 | -0.453 | 0.646 | 0.34 | 0.731 | 1.000 |
| Paraburkholderia | K | Comamonas | Ba | Bb | 0.688 | 0.255 | 0.189 | 1.187 | 2.70 | 0.007 | 0.041 |
| Paraburkholderia | K | Comamonas | Ba | Bh | 0.573 | 0.223 | 0.136 | 1.010 | 2.57 | 0.010 | 0.051 |
| Paraburkholderia | K | Comamonas | Bb | Bh | -0.115 | 0.229 | -0.564 | 0.335 | -0.50 | 0.616 | 1.000 |
| Paraburkholderia | K | Pseudomonas | Non-AA | Ba | -0.831 | 0.337 | -1.491 | -0.171 | -2.47 | 0.014 | 0.054 |
| Paraburkholderia | K | Pseudomonas | Non-AA | Bb | 0.793 | 0.346 | 0.115 | 1.471 | 2.29 | 0.022 | 0.065 |
| Paraburkholderia | K | Pseudomonas | Non-AA | Bh | 0.536 | 0.314 | -0.079 | 1.151 | 1.71 | 0.088 | 0.175 |
| Paraburkholderia | K | Pseudomonas | Ba | Bb | 1.624 | 0.285 | 1.066 | 2.182 | 5.70 | <0.001 | <0.001 |
| Paraburkholderia | K | Pseudomonas | Ba | Bh | 1.367 | 0.238 | 0.900 | 1.834 | 5.73 | <0.001 | <0.001 |
| Paraburkholderia | K | Pseudomonas | Bb | Bh | -0.257 | 0.257 | -0.760 | 0.246 | -1.00 | 0.316 | 0.316 |
| Paraburkholderia | K | Serratia | Non-AA | Ba | -1.067 | 0.284 | -1.624 | -0.510 | -3.75 | <0.001 | <0.001 |
| Paraburkholderia | K | Serratia | Non-AA | Bb | -0.085 | 0.289 | -0.652 | 0.482 | -0.29 | 0.769 | 0.769 |
| Paraburkholderia | K | Serratia | Non-AA | Bh | -0.315 | 0.267 | -0.839 | 0.209 | -1.18 | 0.238 | 0.714 |
| Paraburkholderia | K | Serratia | Ba | Bb | 0.982 | 0.245 | 0.501 | 1.462 | 4.00 | <0.001 | <0.001 |
| Paraburkholderia | K | Serratia | Ba | Bh | 0.751 | 0.213 | 0.333 | 1.170 | 3.52 | <0.001 | 0.002 |
| Paraburkholderia | K | Serratia | Bb | Bh | -0.230 | 0.224 | -0.669 | 0.209 | -1.03 | 0.304 | 0.714 |
| Paraburkholderia | K | Staphylococcus | Non-AA | Ba | -0.237 | 0.267 | -0.760 | 0.285 | -0.89 | 0.374 | 0.747 |
| Paraburkholderia | K | Staphylococcus | Non-AA | Bb | 0.488 | 0.274 | -0.049 | 1.026 | 1.78 | 0.075 | 0.370 |
| Paraburkholderia | K | Staphylococcus | Non-AA | Bh | 0.100 | 0.255 | -0.400 | 0.599 | 0.39 | 0.696 | 0.747 |
| Paraburkholderia | K | Staphylococcus | Ba | Bb | 0.726 | 0.232 | 0.271 | 1.180 | 3.13 | 0.002 | 0.011 |
| Paraburkholderia | K | Staphylococcus | Ba | Bh | 0.337 | 0.201 | -0.057 | 0.731 | 1.68 | 0.094 | 0.370 |
| Paraburkholderia | K | Staphylococcus | Bb | Bh | -0.389 | 0.217 | -0.815 | 0.038 | -1.79 | 0.074 | 0.370 |
| Paraburkholderia | μmax | Agrobacterium | Non-AA | Ba | -0.153 | 0.174 | -0.493 | 0.187 | -0.88 | 0.377 | 0.753 |
| Paraburkholderia | μmax | Agrobacterium | Non-AA | Bb | 0.415 | 0.186 | 0.049 | 0.780 | 2.23 | 0.026 | 0.130 |
| Paraburkholderia | μmax | Agrobacterium | Non-AA | Bh | 0.071 | 0.176 | -0.274 | 0.416 | 0.40 | 0.687 | 0.753 |
| Paraburkholderia | μmax | Agrobacterium | Ba | Bb | 0.568 | 0.168 | 0.239 | 0.897 | 3.38 | <0.001 | 0.004 |
| Paraburkholderia | μmax | Agrobacterium | Ba | Bh | 0.224 | 0.153 | -0.076 | 0.524 | 1.46 | 0.143 | 0.430 |
| Paraburkholderia | μmax | Agrobacterium | Bb | Bh | -0.344 | 0.172 | -0.681 | -0.007 | -2.00 | 0.045 | 0.182 |
| Paraburkholderia | μmax | Brucella | Non-AA | Ba | -0.158 | 0.156 | -0.464 | 0.147 | -1.01 | 0.310 | 0.339 |
| Paraburkholderia | μmax | Brucella | Non-AA | Bb | 0.474 | 0.158 | 0.165 | 0.783 | 3.00 | 0.003 | 0.011 |

| Response | Trait | Competitor | Group 1 | Group 2 | $\Delta log_2FC$ | SE | Lower 95% CI | Upper 95% CI | z | P | P (Holm) |
|---|---|---|---|---|---|---|---|---|---|---|---|
| Paraburkholderia | μmax | Brucella | Non-AA | Bh | 0.278 | 0.155 | -0.027 | 0.582 | 1.79 | 0.074 | 0.222 |
| Paraburkholderia | μmax | Brucella | Ba | Bb | 0.632 | 0.144 | 0.351 | 0.913 | 4.40 | <0.001 | <0.001 |
| Paraburkholderia | μmax | Brucella | Ba | Bh | 0.436 | 0.136 | 0.169 | 0.703 | 3.21 | 0.001 | 0.007 |
| Paraburkholderia | μmax | Brucella | Bb | Bh | -0.196 | 0.143 | -0.476 | 0.084 | -1.37 | 0.170 | 0.339 |
| Paraburkholderia | μmax | Comamonas | Non-AA | Ba | 0.085 | 0.181 | -0.270 | 0.440 | 0.47 | 0.639 | 1.000 |
| Paraburkholderia | μmax | Comamonas | Non-AA | Bb | 0.352 | 0.187 | -0.015 | 0.719 | 1.88 | 0.060 | 0.360 |
| Paraburkholderia | μmax | Comamonas | Non-AA | Bh | 0.226 | 0.180 | -0.127 | 0.579 | 1.25 | 0.210 | 0.841 |
| Paraburkholderia | μmax | Comamonas | Ba | Bb | 0.267 | 0.170 | -0.066 | 0.600 | 1.57 | 0.116 | 0.578 |
| Paraburkholderia | μmax | Comamonas | Ba | Bh | 0.141 | 0.158 | -0.169 | 0.450 | 0.89 | 0.372 | 1.000 |
| Paraburkholderia | μmax | Comamonas | Bb | Bh | -0.126 | 0.168 | -0.455 | 0.203 | -0.75 | 0.451 | 1.000 |
| Paraburkholderia | μmax | Pseudomonas | Non-AA | Ba | -0.255 | 0.266 | -0.776 | 0.265 | -0.96 | 0.336 | 0.336 |
| Paraburkholderia | μmax | Pseudomonas | Non-AA | Bb | 1.559 | 0.304 | 0.964 | 2.155 | 5.13 | <0.001 | <0.001 |
| Paraburkholderia | μmax | Pseudomonas | Non-AA | Bh | 0.830 | 0.267 | 0.308 | 1.353 | 3.11 | 0.002 | 0.006 |
| Paraburkholderia | μmax | Pseudomonas | Ba | Bb | 1.815 | 0.276 | 1.273 | 2.356 | 6.57 | <0.001 | <0.001 |
| Paraburkholderia | μmax | Pseudomonas | Ba | Bh | 1.086 | 0.231 | 0.632 | 1.539 | 4.69 | <0.001 | <0.001 |
| Paraburkholderia | μmax | Pseudomonas | Bb | Bh | -0.729 | 0.277 | -1.272 | -0.186 | -2.63 | 0.008 | 0.017 |
| Paraburkholderia | μmax | Serratia | Non-AA | Ba | -0.364 | 0.252 | -0.857 | 0.129 | -1.45 | 0.148 | 0.593 |
| Paraburkholderia | μmax | Serratia | Non-AA | Bb | 0.322 | 0.278 | -0.222 | 0.866 | 1.16 | 0.246 | 0.739 |
| Paraburkholderia | μmax | Serratia | Non-AA | Bh | 0.068 | 0.250 | -0.423 | 0.559 | 0.27 | 0.787 | 0.787 |
| Paraburkholderia | μmax | Serratia | Ba | Bb | 0.686 | 0.254 | 0.188 | 1.184 | 2.70 | 0.007 | 0.042 |
| Paraburkholderia | μmax | Serratia | Ba | Bh | 0.432 | 0.221 | -0.001 | 0.865 | 1.95 | 0.051 | 0.254 |
| Paraburkholderia | μmax | Serratia | Bb | Bh | -0.254 | 0.253 | -0.749 | 0.241 | -1.01 | 0.314 | 0.739 |
| Paraburkholderia | μmax | Staphylococcus | Non-AA | Ba | 0.442 | 0.329 | -0.203 | 1.086 | 1.34 | 0.179 | 0.519 |
| Paraburkholderia | μmax | Staphylococcus | Non-AA | Bb | 1.764 | 0.440 | 0.903 | 2.626 | 4.01 | <0.001 | <0.001 |
| Paraburkholderia | μmax | Staphylococcus | Non-AA | Bh | 0.450 | 0.330 | -0.197 | 1.096 | 1.36 | 0.173 | 0.519 |
| Paraburkholderia | μmax | Staphylococcus | Ba | Bb | 1.323 | 0.410 | 0.519 | 2.126 | 3.23 | 0.001 | 0.006 |
| Paraburkholderia | μmax | Staphylococcus | Ba | Bh | 0.008 | 0.287 | -0.554 | 0.570 | 0.03 | 0.978 | 0.978 |
| Paraburkholderia | μmax | Staphylococcus | Bb | Bh | -1.315 | 0.411 | -2.120 | -0.509 | -3.20 | 0.001 | 0.006 |
| Competitor | K | Agrobacterium | Non-AA | Ba | 0.747 | 0.194 | 0.366 | 1.129 | 3.84 | <0.001 | <0.001 |
| Competitor | K | Agrobacterium | Non-AA | Bb | 0.117 | 0.160 | -0.196 | 0.430 | 0.73 | 0.465 | 0.465 |
| Competitor | K | Agrobacterium | Non-AA | Bh | 0.293 | 0.158 | -0.017 | 0.603 | 1.85 | 0.064 | 0.191 |
| Competitor | K | Agrobacterium | Ba | Bb | -0.631 | 0.179 | -0.982 | -0.279 | -3.52 | <0.001 | 0.002 |
| Competitor | K | Agrobacterium | Ba | Bh | -0.454 | 0.176 | -0.798 | -0.110 | -2.59 | 0.010 | 0.039 |
| Competitor | K | Agrobacterium | Bb | Bh | 0.177 | 0.141 | -0.099 | 0.452 | 1.25 | 0.209 | 0.419 |
| Competitor | K | Brucella | Non-AA | Ba | 0.444 | 0.158 | 0.135 | 0.753 | 2.82 | 0.005 | 0.019 |
| Competitor | K | Brucella | Non-AA | Bb | -0.164 | 0.127 | -0.413 | 0.085 | -1.29 | 0.196 | 0.587 |
| Competitor | K | Brucella | Non-AA | Bh | -0.105 | 0.125 | -0.350 | 0.139 | -0.84 | 0.399 | 0.797 |
| Competitor | K | Brucella | Ba | Bb | -0.608 | 0.146 | -0.895 | -0.322 | -4.16 | <0.001 | <0.001 |
| Competitor | K | Brucella | Ba | Bh | -0.549 | 0.140 | -0.824 | -0.275 | -3.92 | <0.001 | <0.001 |
| Competitor | K | Brucella | Bb | Bh | 0.059 | 0.110 | -0.157 | 0.275 | 0.53 | 0.593 | 0.797 |
| Competitor | K | Comamonas | Non-AA | Ba | 0.732 | 0.189 | 0.363 | 1.102 | 3.88 | <0.001 | <0.001 |
| Competitor | K | Comamonas | Non-AA | Bb | 0.351 | 0.147 | 0.063 | 0.639 | 2.39 | 0.017 | 0.068 |
| Competitor | K | Comamonas | Non-AA | Bh | 0.372 | 0.144 | 0.089 | 0.655 | 2.58 | 0.010 | 0.049 |
| Competitor | K | Comamonas | Ba | Bb | -0.381 | 0.175 | -0.725 | -0.038 | -2.18 | 0.029 | 0.088 |
| Competitor | K | Comamonas | Ba | Bh | -0.360 | 0.169 | -0.692 | -0.028 | -2.13 | 0.033 | 0.088 |
| Competitor | K | Comamonas | Bb | Bh | 0.021 | 0.126 | -0.225 | 0.267 | 0.17 | 0.867 | 0.867 |
| Competitor | K | Pseudomonas | Non-AA | Ba | 0.503 | 0.228 | 0.056 | 0.950 | 2.21 | 0.027 | 0.109 |
| Competitor | K | Pseudomonas | Non-AA | Bb | -0.343 | 0.184 | -0.703 | 0.017 | -1.87 | 0.062 | 0.186 |

| Response | Trait | Competitor | Group 1 | Group 2 | $\Delta log_2FC$ | SE | Lower 95% CI | Upper 95% CI | z | P | P (Holm) |
|---|---|---|---|---|---|---|---|---|---|---|---|
| Competitor | K | Pseudomonas | Non-AA | Bh | -0.233 | 0.178 | -0.581 | 0.115 | -1.31 | 0.189 | 0.378 |
| Competitor | K | Pseudomonas | Ba | Bb | -0.846 | 0.215 | -1.267 | -0.425 | -3.94 | <0.001 | <0.001 |
| Competitor | K | Pseudomonas | Ba | Bh | -0.736 | 0.206 | -1.139 | -0.333 | -3.58 | <0.001 | 0.002 |
| Competitor | K | Pseudomonas | Bb | Bh | 0.110 | 0.160 | -0.204 | 0.424 | 0.69 | 0.493 | 0.493 |
| Competitor | K | Serratia | Non-AA | Ba | 0.430 | 0.090 | 0.255 | 0.606 | 4.80 | <0.001 | <0.001 |
| Competitor | K | Serratia | Non-AA | Bb | 0.203 | 0.081 | 0.045 | 0.361 | 2.52 | 0.012 | 0.035 |
| Competitor | K | Serratia | Non-AA | Bh | 0.140 | 0.078 | -0.013 | 0.293 | 1.79 | 0.073 | 0.145 |
| Competitor | K | Serratia | Ba | Bb | -0.227 | 0.081 | -0.386 | -0.068 | -2.80 | 0.005 | 0.021 |
| Competitor | K | Serratia | Ba | Bh | -0.290 | 0.070 | -0.427 | -0.153 | -4.16 | <0.001 | <0.001 |
| Competitor | K | Serratia | Bb | Bh | -0.063 | 0.067 | -0.195 | 0.069 | -0.94 | 0.349 | 0.349 |
| Competitor | K | Staphylococcus | Non-AA | Ba | 0.395 | 0.128 | 0.145 | 0.645 | 3.10 | 0.002 | 0.010 |
| Competitor | K | Staphylococcus | Non-AA | Bb | 0.206 | 0.111 | -0.011 | 0.424 | 1.86 | 0.063 | 0.251 |
| Competitor | K | Staphylococcus | Non-AA | Bh | 0.044 | 0.106 | -0.165 | 0.252 | 0.41 | 0.680 | 0.680 |
| Competitor | K | Staphylococcus | Ba | Bb | -0.189 | 0.118 | -0.421 | 0.043 | -1.60 | 0.110 | 0.266 |
| Competitor | K | Staphylococcus | Ba | Bh | -0.351 | 0.108 | -0.562 | -0.140 | -3.26 | 0.001 | 0.007 |
| Competitor | K | Staphylococcus | Bb | Bh | -0.162 | 0.095 | -0.349 | 0.025 | -1.70 | 0.089 | 0.266 |
| Competitor | μmax | Agrobacterium | Non-AA | Ba | 0.266 | 0.238 | -0.201 | 0.733 | 1.12 | 0.264 | 0.528 |
| Competitor | μmax | Agrobacterium | Non-AA | Bb | -0.655 | 0.275 | -1.193 | -0.116 | -2.38 | 0.017 | 0.086 |
| Competitor | μmax | Agrobacterium | Non-AA | Bh | -0.101 | 0.244 | -0.579 | 0.376 | -0.42 | 0.678 | 0.678 |
| Competitor | μmax | Agrobacterium | Ba | Bb | -0.921 | 0.252 | -1.415 | -0.427 | -3.66 | <0.001 | 0.002 |
| Competitor | μmax | Agrobacterium | Ba | Bh | -0.367 | 0.217 | -0.793 | 0.059 | -1.69 | 0.091 | 0.273 |
| Competitor | μmax | Agrobacterium | Bb | Bh | 0.554 | 0.257 | 0.050 | 1.058 | 2.15 | 0.031 | 0.125 |
| Competitor | μmax | Brucella | Non-AA | Ba | 0.290 | 0.083 | 0.128 | 0.452 | 3.50 | <0.001 | 0.002 |
| Competitor | μmax | Brucella | Non-AA | Bb | -0.198 | 0.092 | -0.378 | -0.017 | -2.15 | 0.032 | 0.064 |
| Competitor | μmax | Brucella | Non-AA | Bh | -0.286 | 0.083 | -0.449 | -0.123 | -3.43 | <0.001 | 0.002 |
| Competitor | μmax | Brucella | Ba | Bb | -0.487 | 0.086 | -0.656 | -0.318 | -5.65 | <0.001 | <0.001 |
| Competitor | μmax | Brucella | Ba | Bh | -0.576 | 0.075 | -0.724 | -0.428 | -7.63 | <0.001 | <0.001 |
| Competitor | μmax | Brucella | Bb | Bh | -0.089 | 0.087 | -0.259 | 0.082 | -1.02 | 0.308 | 0.308 |
| Competitor | μmax | Comamonas | Non-AA | Ba | -0.179 | 0.350 | -0.864 | 0.507 | -0.51 | 0.609 | 1.000 |
| Competitor | μmax | Comamonas | Non-AA | Bb | -0.403 | 0.383 | -1.153 | 0.347 | -1.05 | 0.292 | 1.000 |
| Competitor | μmax | Comamonas | Non-AA | Bh | 0.254 | 0.341 | -0.414 | 0.922 | 0.75 | 0.456 | 1.000 |
| Competitor | μmax | Comamonas | Ba | Bb | -0.224 | 0.360 | -0.930 | 0.482 | -0.62 | 0.534 | 1.000 |
| Competitor | μmax | Comamonas | Ba | Bh | 0.433 | 0.315 | -0.185 | 1.051 | 1.37 | 0.170 | 0.848 |
| Competitor | μmax | Comamonas | Bb | Bh | 0.657 | 0.352 | -0.032 | 1.346 | 1.87 | 0.062 | 0.370 |
| Competitor | μmax | Pseudomonas | Non-AA | Ba | -0.124 | 0.076 | -0.273 | 0.024 | -1.64 | 0.100 | 0.500 |
| Competitor | μmax | Pseudomonas | Non-AA | Bb | -0.096 | 0.085 | -0.263 | 0.071 | -1.12 | 0.261 | 1.000 |
| Competitor | μmax | Pseudomonas | Non-AA | Bh | -0.187 | 0.076 | -0.337 | -0.038 | -2.46 | 0.014 | 0.083 |
| Competitor | μmax | Pseudomonas | Ba | Bb | 0.028 | 0.079 | -0.126 | 0.183 | 0.36 | 0.719 | 1.000 |
| Competitor | μmax | Pseudomonas | Ba | Bh | -0.063 | 0.067 | -0.194 | 0.068 | -0.94 | 0.346 | 1.000 |
| Competitor | μmax | Pseudomonas | Bb | Bh | -0.091 | 0.079 | -0.247 | 0.064 | -1.15 | 0.250 | 1.000 |
| Competitor | μmax | Serratia | Non-AA | Ba | 0.336 | 0.051 | 0.235 | 0.437 | 6.54 | <0.001 | <0.001 |
| Competitor | μmax | Serratia | Non-AA | Bb | 0.124 | 0.056 | 0.014 | 0.235 | 2.21 | 0.027 | 0.054 |
| Competitor | μmax | Serratia | Non-AA | Bh | 0.132 | 0.051 | 0.033 | 0.232 | 2.60 | 0.009 | 0.028 |
| Competitor | μmax | Serratia | Ba | Bb | -0.212 | 0.053 | -0.315 | -0.108 | -4.02 | <0.001 | <0.001 |
| Competitor | μmax | Serratia | Ba | Bh | -0.204 | 0.044 | -0.290 | -0.118 | -4.64 | <0.001 | <0.001 |
| Competitor | μmax | Serratia | Bb | Bh | 0.008 | 0.052 | -0.094 | 0.109 | 0.15 | 0.882 | 0.882 |
| Competitor | μmax | Staphylococcus | Non-AA | Ba | 0.027 | 0.039 | -0.049 | 0.103 | 0.69 | 0.488 | 0.976 |
| Competitor | μmax | Staphylococcus | Non-AA | Bb | -0.129 | 0.044 | -0.215 | -0.043 | -2.94 | 0.003 | 0.010 |

| Response | Trait | Competitor | Group 1 | Group 2 | $\Delta log_2FC$ | SE | Lower 95% CI | Upper 95% CI | z | P | P (Holm) |
|---|---|---|---|---|---|---|---|---|---|---|---|
| Competitor | µmax | Staphylococcus | Non-AA | Bh | -0.136 | 0.040 | -0.214 | -0.058 | -3.42 | <0.001 | 0.002 |
| Competitor | µmax | Staphylococcus | Ba | Bb | -0.156 | 0.040 | -0.234 | -0.078 | -3.94 | <0.001 | <0.001 |
| Competitor | µmax | Staphylococcus | Ba | Bh | -0.163 | 0.030 | -0.222 | -0.103 | -5.34 | <0.001 | <0.001 |
| Competitor | µmax | Staphylococcus | Bb | Bh | -0.007 | 0.040 | -0.086 | 0.072 | -0.17 | 0.868 | 0.976 |
