## Supplemental Figures for "*Paraburkholderia* endosymbionts with streamlined genomes are disadvantaged during interspecific competition"

### Supplementary Figures

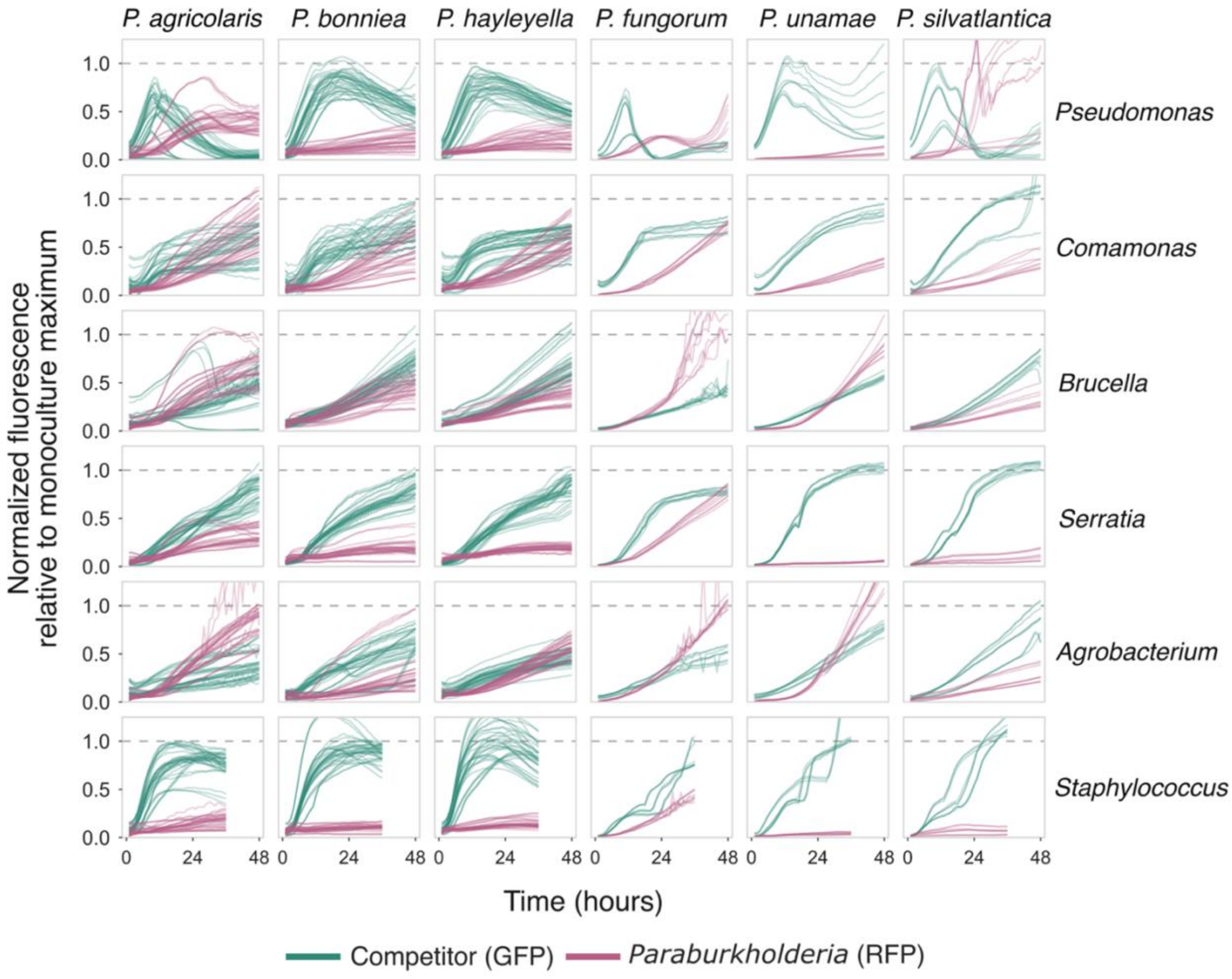


**Figure S1: Monoculture-normalised fluorescence time curves across competitive co-cultures.** Bacterial pairs were inoculated into liquid yeast-glucose (YG) medium at equal starting ratios. RFP and GFP fluorescence were measured hourly for 48 h using a microplate reader. Green curves represent GFP-labelled competitor strains (rows), and pink curves represent RFP-labelled *Paraburkholderia* strains (columns). Each fluorescence curve was normalised to the corresponding strain's maximum fluorescence in same-day monoculture, following correction for background fluorescence and cross-channel bleed-through. The dashed horizontal line indicates the monoculture maximum. Individual curves represent technical replicates across experimental dates, which were averaged within dates for downstream analyses. *Staphylococcus* co-cultures were truncated at 36 h due to atypical late-stage fluorescence dynamics.

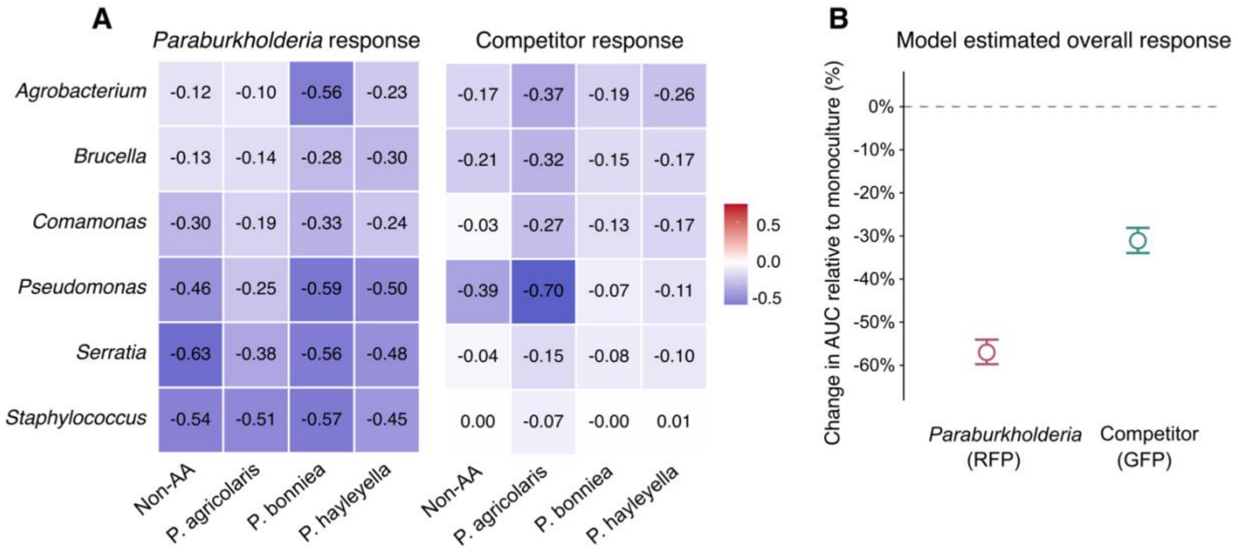


**Figure S2: Effects of interspecific competition on monoculture-normalised fluorescence AUC. (A)** Heatmaps show mean $\log_{10}$-transformed changes in fluorescence AUC relative to same-day monocultures, following correction for background fluorescence and cross-channel bleed-through. Values were averaged across experimental dates and strains within each *Paraburkholderia* group. The left heatmap shows the *Paraburkholderia* response (RFP), and the right shows the competitor response (GFP). Cooler colours indicate greater reductions in AUC, whereas warmer colours indicate increases. **(B)** Model-estimated percentage changes in AUC relative to monoculture, averaged equally across six *Paraburkholderia* species and six competitors. Points show back-transformed estimated marginal means with 95% confidence intervals. Competition reduced AUC for both *Paraburkholderia* (57.0%) and competitors (30.8%; both Holm-adjusted $p < 0.001$).

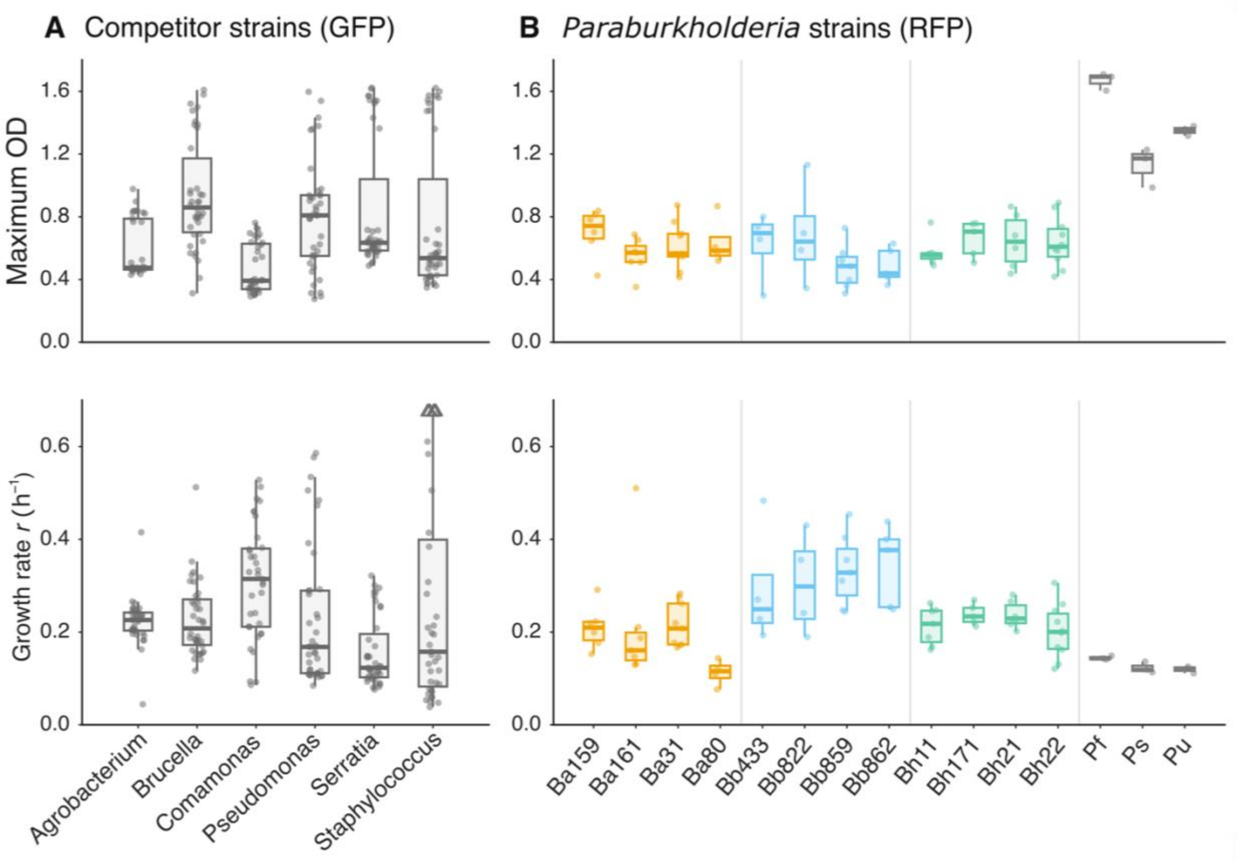


**Figure S3: Optical density ($OD_{600}$)-derived growth characteristics of bacterial strains in monoculture.** Maximum $OD_{600}$ was measured over 0–48 h, and maximum growth rate ($\mu_{max}$) was estimated over the first 24 h. **(A)** GFP-labelled competitor strains and **(B)** RFP-labelled *Paraburkholderia* strains. Upper panels show maximum $OD_{600}$, and lower panels show μmax. Boxplots summarise the distributions of measurements, with points representing individual observations. Colours distinguish *P. agricolaris* (orange), *P. bonniea* (blue), *P. hayleyella* (green), and the non-amoeba-associated species (grey).

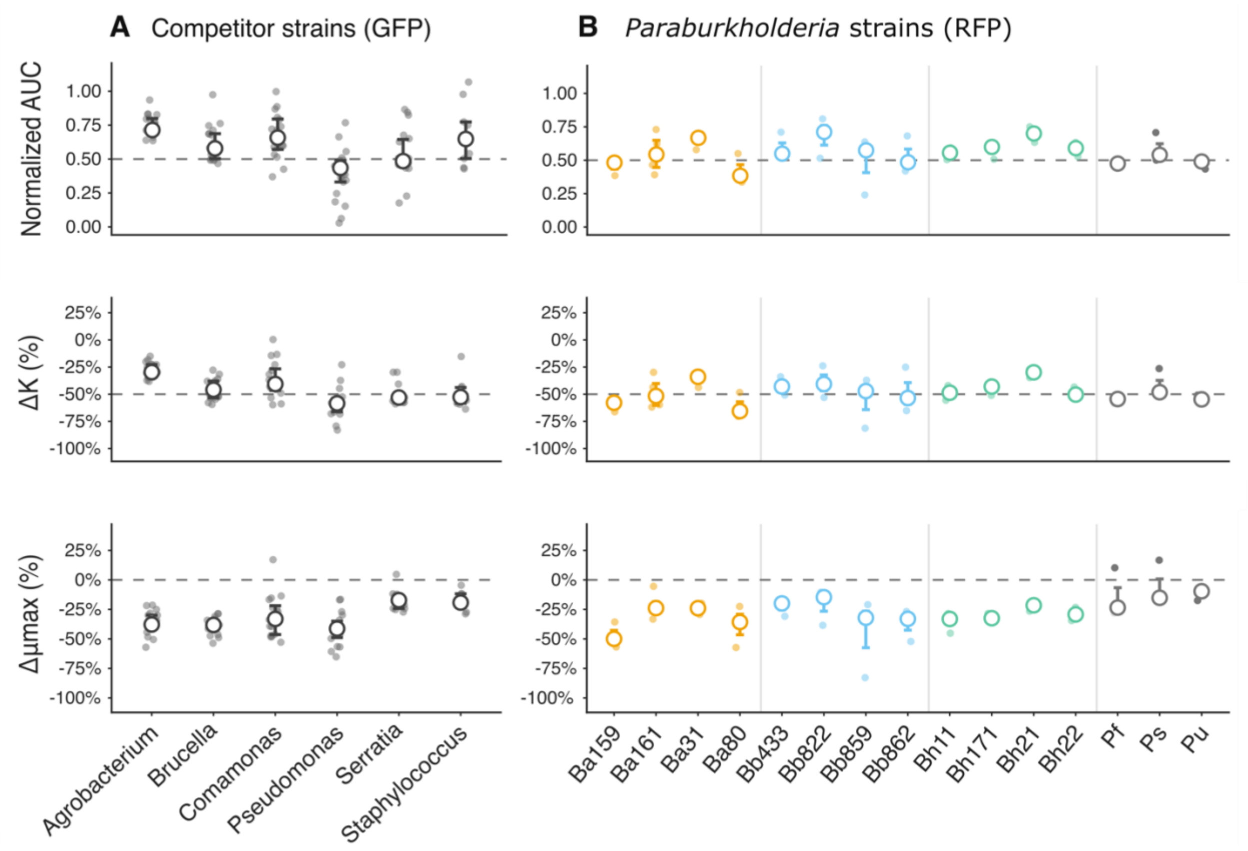


**Figure S4: Growth of fluorescently labelled strains with matched unlabelled wild-type controls.** Fluorescent strains were grown at equal 1:1 starting ratios with their corresponding unlabelled wild-type (WT) strains. **(A)** GFP-labelled competitor strains and **(B)** RFP-labelled *Paraburkholderia* strains. Rows show monoculture-normalised fluorescence AUC (top), percentage change in maximum fluorescence yield (*K*; middle), and percentage change in maximum growth rate (μmax; bottom), relative to same-day monocultures. Small points represent experimental-date observations, while open circles show strain medians with interquartile ranges. Dashed lines indicate theoretical expectations under equal resource sharing: 0.5 for normalised AUC, −50% for *K*, and 0% for μmax. *Paraburkholderia* strains are coloured by group: *P. agricolaris* (orange), *P. bonniea* (blue), *P. hayleyella* (green), and non-amoeba-associated species (grey).

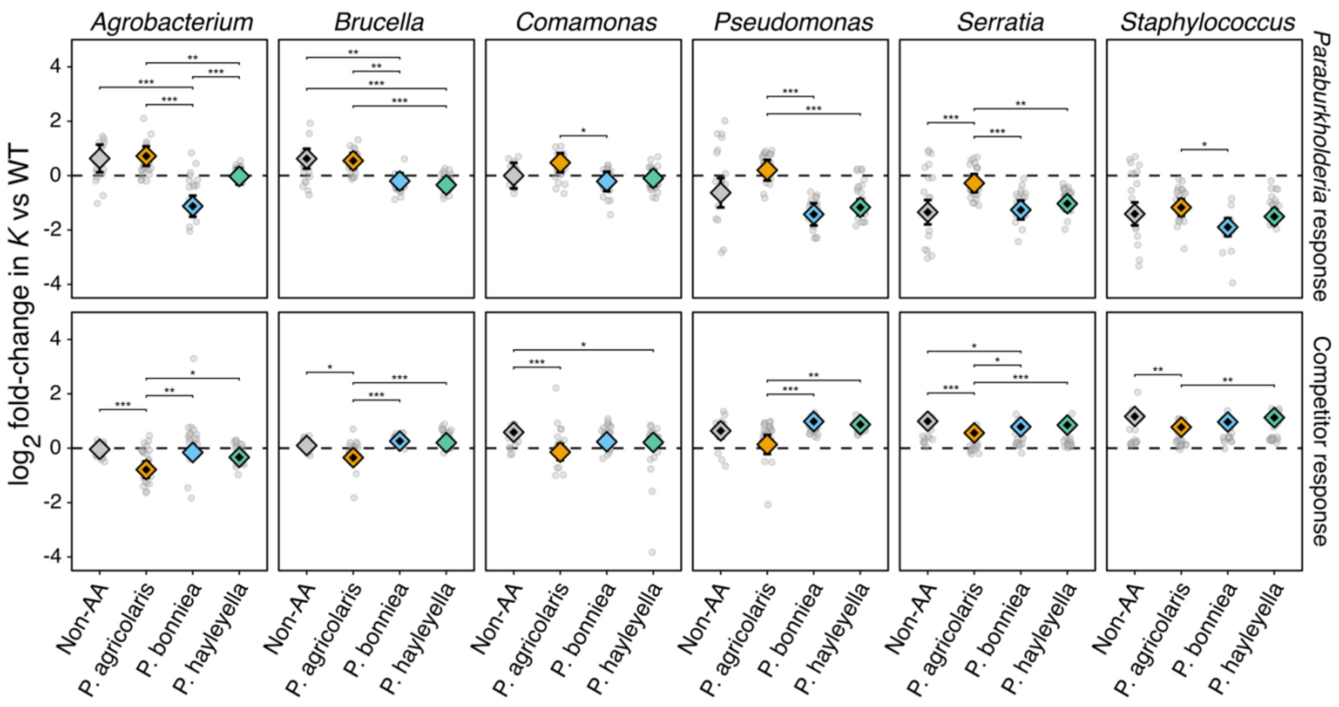


**Figure S5: Competitor-specific fluorescence-yield responses relative to WT.** Maximum fluorescence yield (*K*) is expressed as $\log_2$ fold-change relative to matched labelled–unlabelled wild-type (WT) co-culture controls, with zero indicating no deviation from this baseline. Columns show competitor species, and rows separate *Paraburkholderia* (upper) and competitor (lower) responses. Grey points represent experimental-date observations, while coloured diamonds show model-estimated marginal means with 95% confidence intervals. Black centres within diamonds indicate significant departures from the WT baseline, with Holm adjustment across 24 tests within each response. Significance brackets indicate planned pairwise contrasts among *Paraburkholderia* groups, with Holm adjustment across six comparisons within each competitor × response (* $p < 0.05$; ** $p < 0.01$; *** $p < 0.001$).

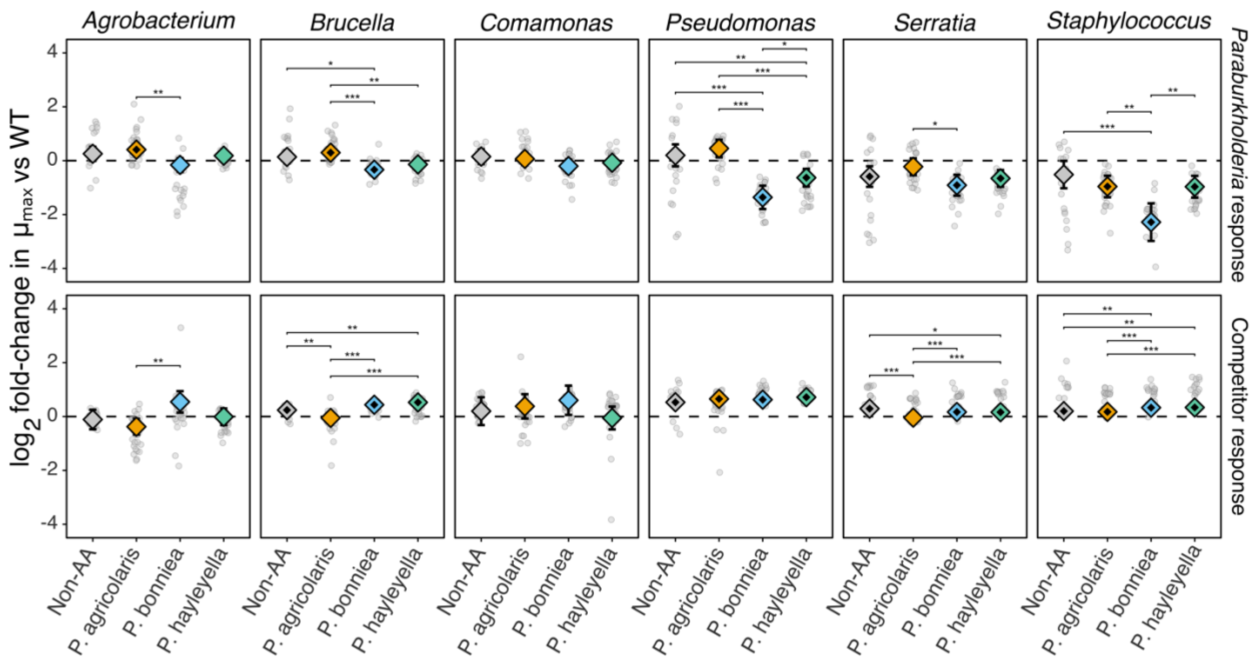


**Figure S6: Competitor-specific maximum growth-rate responses relative to WT.** Fluorescence-derived maximum growth rate ($\mu_{max}$) is expressed as $log_2$ fold-change relative to matched labelled–unlabelled wild-type (WT) co-culture controls, with zero indicating no deviation from this baseline. Columns show competitor species, and rows separate *Paraburkholderia* (upper) and competitor (lower) responses. Grey points represent experimental-date observations, while coloured diamonds show model-estimated marginal means with 95% confidence intervals. Black centres within diamonds indicate significant departures from the WT baseline, with Holm adjustment across 24 tests within each response. Significance brackets indicate planned pairwise contrasts among *Paraburkholderia* groups, with Holm adjustment across six comparisons within each competitor × response (* $p < 0.05$; ** $p < 0.01$; *** $p < 0.001$).

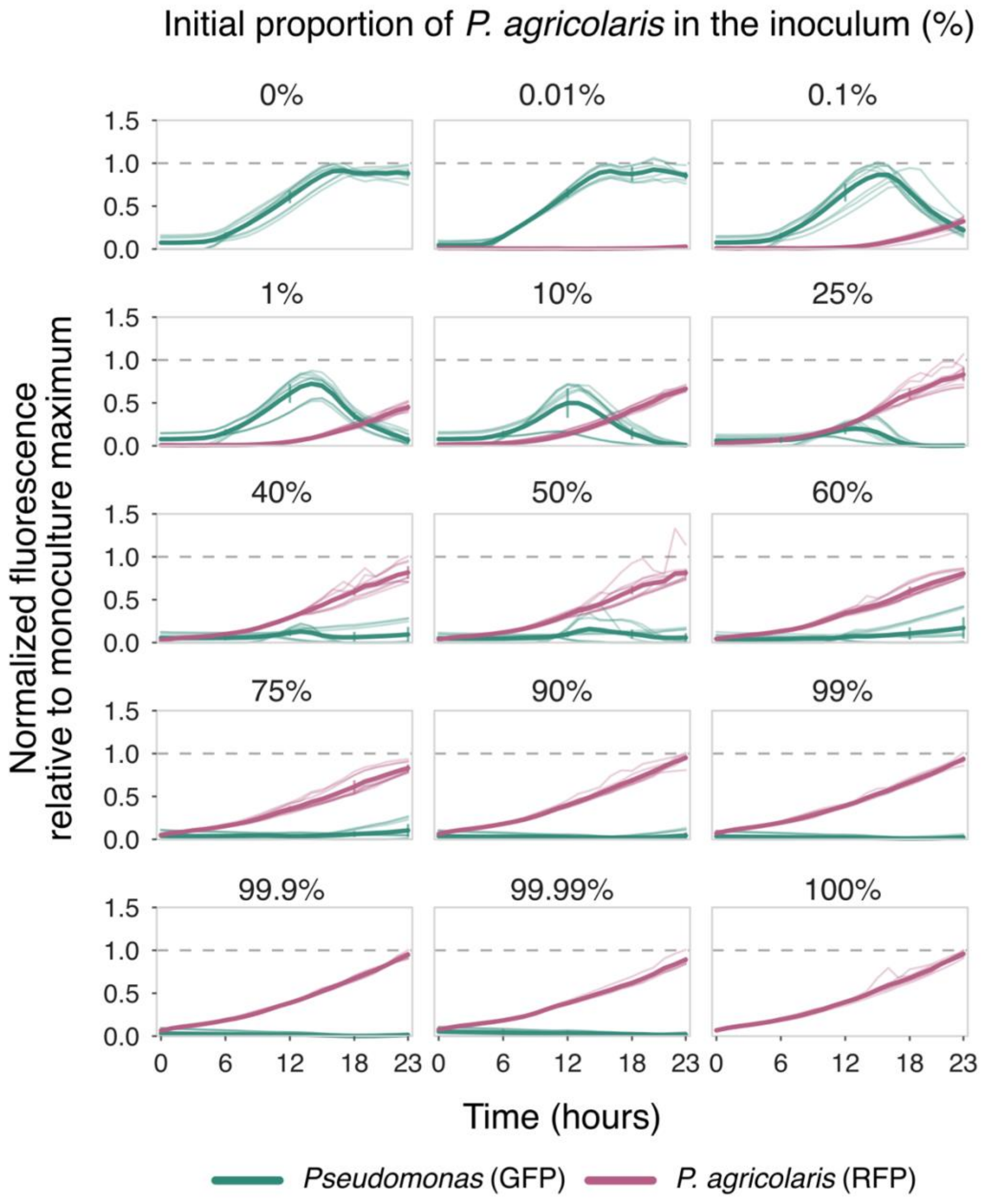


**Figure S7: Fluorescence dynamics across varying starting ratios of *P. agricolaris* and *Pseudomonas***. *P. agricolaris* strain Ba161 and *Pseudomonas* were co-cultured at 15 initial Ba161 proportions (panel headings), with fluorescence monitored over 24 h. Green curves represent GFP-labelled *Pseudomonas*, and pink curves represent RFP-labelled *P. agricolaris*. Fluorescence was normalised to the corresponding same-day monoculture maximum, with the dashed line indicating a value of 1. Individual curves show replicate trajectories. *Pseudomonas* fluorescence declined following initial growth even when Ba161 comprised only 0.1% of the starting inoculum**.**
